# Measurement of a panel of 21 steroids in a quantitative assay in human plasma, adipose tissue, and fecal samples using ultra-high-performance liquid chromatography–tandem mass spectrometry

**DOI:** 10.64898/2026.07.08.737297

**Authors:** Ilia Evstafev, Matilda Kråkström, Niina Saarinen-Aaltonen, Janne Hakkarainen, Merja R. Häkkinen, Seppo Auriola, Peter J. Boström, Matti Poutanen, Matej Orešič, Alex M. Dickens

## Abstract

Comprehensive detection of steroids, beyond the limited panels typically analyzed in clinical chemistry laboratories, has become increasingly important given their pivotal roles in diverse biological processes. However, steroid quantification poses several analytical challenges, including differences in ionization efficiency and structural similarities across the entire steroid metabolic network. To address these challenges, we developed a targeted ultra-high-performance liquid chromatography–tandem mass spectrometry (UHPLC–MS/MS) assay to analyze 21 steroids using reverse-phase chromatography combined with rapid polarity switching. Mass spectrometry (MS) analysis was performed in scheduled multiple reaction monitoring (sMRM) mode. Depending on the steroid and matrix, the validated lower limits of quantitation (LLOQ) ranged from 12.0 pM to 1216 pM in plasma and 41.1 pM to 384 pM in fecal sample homogenates. In adipose tissue, it was from 0.01 pmol/g to 9 pmol/g. Measured steroid concentrations obtained from the commercial control samples (MassTrak™ Steroid Serum QC Set 1 and the MassCheck® Steroid Panel 1 Serum Control) showed close agreement with the reference values. As a proof of concept, the method was successfully applied to 469 plasma samples in several projects, 15 adipose tissue samples, and 332 fecal samples, demonstrating its applicability to large-scale studies. In conclusion, the method enables sensitive, derivatization-free quantification of an expanded steroid panel in plasma and complex biological matrices, including adipose tissue and fecal samples, representing a significant advancement in comprehensive steroid profiling.

**Graphical abstract:** For Table of Contents Only

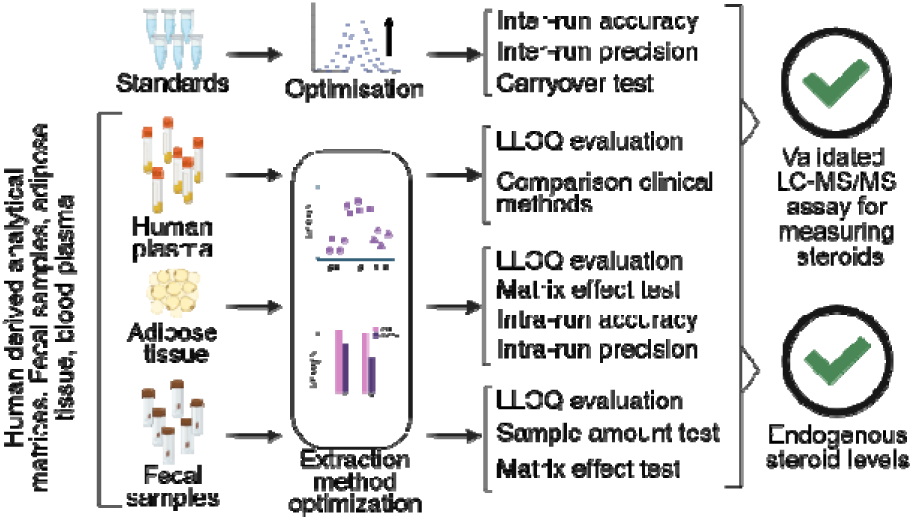

---

Steroid hormones, including mineralocorticoids, glucocorticoids, and sex steroids, play central roles in regulating energy and electrolyte balance, immunity, cardiovascular function, and reproduction in humans and across a variety of animal species^1, 2^. Because human and animal studies often rely on limited sample volumes, highly sensitive analytical methods are required for accurate quantification of steroids^2^. This need is further amplified in animal studies, such as mice, where steroid concentrations are typically lower than in humans, ranging from pM to nM concentrations^3^. Consequently, the development of sensitive and robust analytical techniques is crucial for reliable steroid detection and quantification across species.

Given the fundamental importance of steroids in biological systems, the need for accurate analytical methods has long been recognized. Immunoassays are still the most widely applied approaches for steroid measurement, having been in use for over 50 years^4^. These were initially designed for the detection of protein hormones^5^, and were soon adapted for steroid analysis^6^. However, numerous studies have reported that even modern immunoassays can yield biased results due to limited specificity and selectivity, particularly when compared with mass spectrometry-based methods such as liquid chromatography–mass spectrometry (LC-MS) and gas chromatography–mass spectrometry (GC-MS)^7, 8^. Additionally, immunoassays are generally designed to quantify a single analyte, and expansion to additional targets is labor-intensive, requiring the development of new specific antibodies and additional sample volume^4^. As a result, the selection of analytes in clinical practice is often restricted to a small subset of steroids, limiting the ability to interrogate less abundant or pathway-specific components within complex biological matrices^4^. Furthermore, because steroids are poorly immunogenic, many immunoassays lack the optimal sensitivity required for animal studies^9^. In contrast, MS enables multiplexed analysis without such constraints^10^, and thus, in recent years, interest in broad steroid profiling of tissue samples has increased, driven also by the increased sensitivity of MS.

Established GC-MS and LC-MS methods have demonstrated robust performance for profiling a wide range of steroids in human serum and plasma^11–13^. Traditionally, derivatization has been used to increase the sensitivity of steroid hormones ^14, 15^, but it adds complexity and limits throughput. For example, a multi-step protocol including homogenization, ethyl acetate extraction, several derivatization steps and SPE separation steps was validated for the determination of 8 steroid species in human adipose tissue, including cortisol, estradiol, and testosterone, with 5-300 pg/g LLOQs^16^. These studies confirmed that adipose tissue actively participates in steroid hormone biosynthesis and functions as an endocrine organ. However, other sample types, such as fecal matter, have received less attention, and such studies have been mainly carried out on animals^17–19^.

Adipose tissue represents a major reservoir for steroid hormones both in human and in mouse models due to their lipophilic nature^20, 21^, expresses estrogen, androgen, and glucocorticoid steroid receptors^22^ and the enzymatic machinery required for steroid metabolism^23^, highlighting its role in the establishment of endocrine homeostasis and underscoring its relevance for comprehensive pathway analysis. Despite this, various intermediates of steroid synthesis and metabolism are rarely interrogated. Another growing area of study is the gut microbiome, and several studies have pointed to this also being an endocrine organ with the bacteria expressing an enzyme machinery to metabolise steroids^24–34^. Furthermore, data have shown an association between circulating sex steroid levels and the composition of the gut microbiota in both humans and preclinical animal models^25, 29, 33, 35^. Yet, so far, human studies remain largely restricted to narrow steroid panels. Together, the gaps described above highlight the need for validated analytical methods capable of quantifying expanded steroid panels in both adipose tissue and fecal samples.

In this study, we describe the development of a targeted ultra-performance liquid chromatography–tandem mass spectrometry (UHPLC-MS/MS) assay for quantifying 21 steroids, including the active hormones and various precursor steroids. Importantly, the method does not require derivatization while achieving detection limits comparable to those of derivatization-based approaches^36, 37^. The method is expected to have a wide range of applications in basic and clinical research, and to produce novel data that can be used to interpret disease mechanisms and to diagnose and prevent them.

## MATERIALS AND METHODS

### REAGENTS AND STANDARDS

Water, methanol (MeOH), acetonitrile (MeCN) (all, CHROMASOLV™ LC-MS Ultra); tert-butyl methyl ether (MTBE) HPLC grade was obtained from Honeywell (Riedel-de Haën Charlotte, North Carolina, USA). MassTrak Steroid Serum QC Set 1 was acquired from Waters (Milford, MA, USA). Pooled human plasma was obtained from Biowest (Nuaille, France). 11β-OH-androstenedione (11β-OHA4), 11-keto-dihydrotestosterone (11-KDHT), and 11-ketotestosterone (11-KT) were obtained from Cayman (Chemical Ann Arbor, MI, USA); androsterone (AN) and pregnenolone (P5) were obtained from Fisher Scientific (Waltham, MA, USA); 11-deoxycorticosterone (DOC), 17α-OH-progesterone (17α-OHP4), adrenosterone (11-KA4), aldosterone (A), androstenedione (A4), corticosterone (B), cortisol (F), cortisone (E), dehydroepiandrosterone (DHEA), dihydrotestosterone (DHT), estradiol (E2), estrone (E1), progesterone (P4), and testosterone (T) were obtained from Sigma-Aldrich (Saint Louis, MO, USA); 11-deoxycortisol (S) and 17α-OH-pregnenolone (17α-OHP5) were obtained from Toronto Research Chemicals (North York, ON, Canada). Internal standards 11-ketotestosterone-d3 (d3-11-KT), and 5α-dihydro-11-ketotestosterone-d3 (d3-11-KDHT) were obtained from Cayman Chemical (Ann Arbor, MI, USA); testosterone-d3 (d3-T), aldosterone-d7 (d7-A), and dihydrotestosterone-d4 (d4-DHT) from IsoSciences (Ambler, PA, USA); androsterone-d4 (d4-AN), and 11β-OH-4-androstene-3,17-dione-d4 (d4-11β-OHA4) were obtained from Sigma-Aldrich (Saint Louis, MO, USA); 17α-hydroxy-pregnenolone-d3 (d3-17α-OHP5), corticosterone-d8 (d8-B), pregnenolone-d4 (d4-P5), 11-deoxy-cortisol-d7 (d7-S), 11-deoxy-corticosterone-d7 (d7-DOC), cortisol-d4 (d4-F), progesterone-d9 (d9-P4), cortisone-d8 (d8-E), 17α-hydroxy-progesterone-d8 (d8-17α-OHP4), 17β-estradiol-d4 (d4-E2), estrone-d4 (d4-E1), dehydroepiandrosterone-d2 (d2-DHEA), and androstenedione-d3 (d3-A4) were obtained from Toronto Research Chemicals (North York, ON, Canada). CAS numbers of analytes could be found in supplementary materials (Table S1).

### HUMAN SAMPLES

Commercially available flash-frozen subcutaneous human adipose tissue samples were originally derived from three healthy premenopausal female donors under informed patient consent and with the approval of Institutional Review Board (consent and approvals by Zenbio, Inc.) and used for method validation and development. Commercially available pooled human plasma collected using citrate-phosphate-dextrose anticoagulant was used for the test protocols (S4180, Biowest, Nuaille, France). The fecal samples were obtained as part of the PROMIC study (https://clinicaltrials.gov/study/NCT06116851) and were collected with OMNImet•GUT kit, DNA (Genotek Ontario, Canada).

### PREPARATION OF STANDARD SOLUTIONS

Internal standard (IS) solution consisting of deuterium-labeled steroids was prepared in MeCN:water mixture (30:70, v:v). The concentrations were optimized to achieve a proper response during the analysis without introducing noticeable amounts of native compounds as impurities and preventing their interference with the analysis. The concentrations were selected as follows: 2 nM for d3-11-KDHT, and d3-A4; 4 nM for d3-T; 20 nM for d8-B, d7-S, d7-DOC, d4-F, d9-P4, d8-E, d8-17α-OHP4, d7-A, d3-11-KT, and d4-11β-OHA4; 40 nM for d4-E2, d4-E1, and d4-AN; 100 nM for d4-DHT; 200 nM for d3-17α-OHP5, d4-P5, and d2-DHEA. Working solutions, comprised of native steroid standards were prepared in a mixture of MeCN:water (30:70, v:v) at 16 concentration levels by serial dilution from 0.005 nM to 1000 nM (Table S2). This allowed us to fully define the dynamic range for the selected analytes. For routine analyses, we modified the standard curve to a 10-point calibration curve prepared from working solutions ranging from 0.25 nM to 1000 nM. The calibration curve samples were prepared by mixing 20 µL of the IS solution, 20 µL of the corresponding working solution, and 10 µL of MeCN:water mixture (30:70, v:v). Solvent quality controls (QCs) were used to verify method performance during the analysis. The Solvent QC samples were prepared using the same procedure as for the calibration curve sample

### SERUM AND PLASMA LIQUID-LIQUID EXTRACTION (LLE) PROCEDURE

A 150 µL volume of human plasma or serum was spiked with 20 µL of IS solution and vortexed. One mL of MTBE was added to the obtained mixture and vortexed for 10 min. A 900 µL volume from the top MTBE layer was withdrawn, filtered through a protein precipitation plate (55263-U Supelco (PA, USA), transferred into total recovery vials (Waters 186002805, Milford, MA, USA), and evaporated. The samples were reconstituted in 50 µL of MeCN:water (30:70, v:v)

### SERUM AND PLASMA PROTEIN PRECIPITATION (PP) PROCEDURE

Sample preparation was carried out in accordance with the abovementioned LLE extraction, except 500 µl of cooled methanol was used instead of 1 ml of MTBE.

### SERUM AND PLASMA SUPPORTED LIQUID EXTRACTION (SLE) PROCEDURE

The extraction setup was based on the technical note TN78770221_W, Phenomenex (Torrance, California). Samples were pre-treated by adding 200 µL of phosphate-buffered saline to 150 µL of human plasma and 20 µL of IS solution. Pre-treated samples were loaded onto a Novum SLE PRO 96-Well Plate, Phenomenex (Torrance, California), in order to extract steroids. A vacuum was applied until the samples had completely entered the sorbent. The elution was carried out with 2× 900 µL of ethyl acetate. A vacuum of approximately 2 minutes was applied. The eluate was evaporated to dryness at 40 °C for 15 min and reconstituted in 50 µL of MeCN:water (30:70, v:v)

### ADIPOSE TISSUE TWO-STEP LLE (2LLE) PREPARATION

Adipose tissue (approximately, but not less than 20 mg) was weighed and homogenized in a MeOH:water (1:1, v:v) solution at a ratio of 1:5 (w:v). The samples were kept on dry ice, and homogenization was carried out using a bullet blender (Next Advance Troy, NY, USA), with 0,5 mm diameter zirconium oxide beads, ZROB05 (Next Advance, Troy, NY, USA) at 2-8 °C for two periods of 1 minute with 7/10 speed settings, then samples were returned on dry ice until the extraction. Adipose tissue homogenate (120 µL) was spiked with 20 µL of IS solution and vortexed. Hexane (800 μL) was added, and the tube was vortexed thoroughly. Then 800 µL MeOH:water 1:1 (v:v) was added, vortexed, centrifuged, and the MeOH:water (bottom) layer was collected. Another aliquot (800 µL) of MeOH:water 1:1 (v:v) was added, the mixture vortexed, centrifuged, and the MeOH:water (bottom) layer was collected. The two collected fractions were combined and dried, and the residue was reconstituted in 50 µL of MeCN: water mixture (30:70, v:v).

### FECAL SAMPLE PREPARATION PROCEDURE

Each sample was split into three aliquots. A mixture of EtOH:Water (1:1 v:v) was added to a sample aliquot to bring the final volume to 1-1.25 mL. Zirconium oxide beads were added to each tube, then all the samples underwent homogenization in a bullet blender, (Next Advance, Troy, NY, USA) for 2 minutes (with 10/12 speed settings). A 100 µL aliquot of the resulting slurry was spiked with 20 µL of the IS solution and vortexed. One mL of MTBE was added to the resultant mixture and vortexed for 10 min. The mixture was stored in -80°C overnight. When the lower layer was frozen, the samples were centrifuged at 14 000 rpm for 10 minutes. Then, 900 µL was withdrawn, filtered through a protein precipitation plate (Product code: 55263-U Supelco PA, USA), transferred to total recovery vials (Product code:186002805 Waters Milford, MA, USA), evaporated, and reconstituted in 50 µL of MeCN:water (30:70, v:v). To avoid injection of clogged lipid samples, total recovery vials were centrifuged at 1000 rpm for 5 minutes. An aliquot (100 μL) of the fecal slurry was subjected to a 24-hour drying cycle using a CentriVap, (Labconco Kansas City, MO, USA). The process included 21 hours of drying at room temperature followed by a further 3 hours at 35°C. Dry matter content was determined as the difference between the weight of the empty tube and its weight after drying.

### SELECTION OF EXTRACTION PROCEDURE IN PLASMA

To compare plasma extraction procedures, pooled plasma samples were prepared in triplicate using LLE, PP, and SLE protocols that were modified accordingly. The samples were spiked with a working solution of native steroids (50 nM) to mimic samples containing elevated analyte concentrations. The IS solution was added after extraction. Additionally, a set of samples (n=3) was prepared in neat solutions with target concentrations. All samples were analyzed against calibration curves.

### SELECTION OF ADIPOSE TISSUE EXTRACTION PROCEDURE

To study the distribution of analytes between extraction phases during the 2LLE procedure, a set of adipose tissue samples (n = 3) was prepared. In this set, both the hexane and MeOH:water layers were collected and analyzed, and the second extraction step was omitted.

For evaluation of extraction efficiency, two sample sets were prepared in triplicate from adipose tissue using the 2LLE protocol, with one set including the second extraction step and the other omitting it. The IS solution was added at the final stage of these sample preparation to account for losses of native steroids during extraction. Additionally, IS compensates for matrix effects and LC-MS system variability. The prepared test samples were analyzed in one batch and quantified using a calibration curve prepared in solvent.

### LLOQ EVALUATION

The endogenous nature of steroid hormones makes it nearly impossible to obtain a blank matrix and use it to determine the LLOQs^38^. For plasma and fecal samples, we assessed the LLOQ in matrix by normalization on the IS response. A conversion coefficient was calculated as the ratio of the mean IS response in the matrix to the mean IS response in the neat solution. The LLOQ in neat solutions was defined as the lowest concentration level providing a signal at least 50% higher than that of blank samples, exhibiting reproducible response (CV<20%), and with accuracy within 80–120%. The LLOQs calculated in neat solutions were subsequently divided by this coefficient to estimate the dynamic range in the matrix.

For the adipose tissue samples, we adopted the surrogate analyte approach described in regulatory guidelines^39^. In this approach, we used isotopically labeled steroids as surrogate analytes, with native compounds serving as internal standards. Homogenate aliquots (120 µL) were spiked with isotopically labeled steroids at 16 concentration levels ranging from 0.005 nM to 1000 nM (Table S2) in MeCN:water (30:70, v:v). Subsequently, 20 µL of a working solution of native steroids (100 nM per compound) was added.

The surrogate analyte LLOQ was defined as the lowest concentration providing a signal (AUC) at least 50% higher than that of a blank from the same sample preparation procedure, and a calculated accuracy in the range of 80-120%. The ULOQ was defined as the highest concentration within the linear dynamic range, exhibiting no signs of column overload and an accuracy within 85–115%.

Next, we assessed the conversion of the LLOQ of the surrogate analyte to the LLOQ of the target analyte. We prepared samples (n=3) with identical concentrations (40 nM) of both isotopically labeled and native steroids in MeCN:water (30:70, v:v). We then calculated the signal-to-noise ratios for both the surrogate analytes and the native steroids. The transformation coefficient was calculated as the ratio of the surrogate analyte signal-to-noise to that of the corresponding native analyte. Then, by multiplying by this coefficient, the LLOQ of the isotopically labeled analytes was converted to the LLOQ of the native steroids.

### EVALUATION OF MATRIX EFFECT IN ADIPOSE TISSUE

The matrix effect for adipose tissue was determined by calculating the normalized matrix factor (nMF). To evaluate it, a series of samples from individual subjects (n=6) was prepared according to the above-described two-step LLE procedure, with omission of the IS solution to determine the endogenous signal levels in the test samples. The samples were injected into the UHPLC-MS/MS analysis. Next, the same extracts were spiked with an internal standard and a working solution of known concentration, dried, and brought to the volume remaining after the first injection. The samples were then analyzed together with a corresponding series of matrix-free controls (n=6) prepared in MeCN:water mixture (30:70, v:v). The obtained areas under the curve (AUC) and internal standard areas under the curve (IS AUC) values were used to calculate the normalized matrix effect according to the following formula:

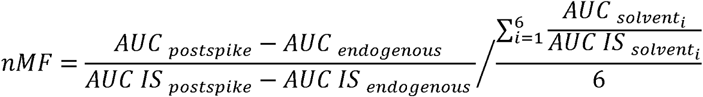

A coefficient of variation (CV) below 15% was used as the acceptance criterion.

### INTRA-RUN REPEATABILITY, PRECISION, AND ACCURACY IN ADIPOSE TISSUE

To assess the performance in the assay in real samples, adipose tissue homogenates (120 µL) were spiked with a working solution of native steroids (20 µL) and IS solution (20 µL). The working solution concentrations were set at the middle of the analytical range: 1 nM for low quality control samples (LQC; equivalent to 1 pmol/g in adipose tissue) and 50 nM for high quality control samples (HQC; equivalent to 50 pmol/g in adipose tissue). Both levels were prepared in 5 replicates using the two-step LLE protocol previously optimized for the adipose tissue. Each replicate was injected 3 times, CV across injections was calculated to assess the repeatability. The intra-run precision was calculated by evaluating CV values across all replicates at the same concentration of spiked samples. A CV threshold of 15% was applied as the acceptance criterion for intra-run repeatability and intra-run precision. The intra-run accuracy was assessed by calculating the accuracy values across all the replicates of the same concentration of spiked samples. The endogenous presence of many of the steroids in the analyzed panel prevented evaluation of accuracy at the LQC level. However, the concentrations measured in the LQC, which reflect the concentration of the steroids in the sample, were used to correct the endogenous steroid level. In order to exclude the endogenous levels of steroids in the matrix and to obtain an accurate assessment of the accuracy of the spiked steroid, the following calculation was performed:

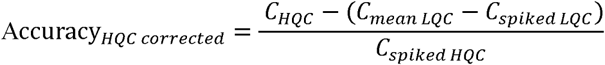

The corrected accuracy was considered acceptable when it was within the 85%-115% range.

### INTER-RUN ACCURACY AND PRECISION USING SOLVENT- AND SERUM-BASED QC SAMPLES

Inter-run accuracy and precision were evaluated using solvent-based QC samples at a concentration of 20 nM. QC samples were analyzed across three independent batches (total n = 39 injections: batch 1, n = 5; batch 2, n = 6; batch 3, n = 28). To assess method comparability in a biological matrix, serum-based QC materials (Waters MassTrak™ and Chromsystems MassCheck® steroid panels) were reconstituted according to the manufacturers’ instructions and subjected to the same LLE extraction procedure as that used for serum and plasma samples. The resulting extracts were analyzed against calibration curves, and concentrations were calculated.

In addition, a cohort of 322 plasma samples from male donors from the PROMIC study (https://clinicaltrials.gov/study/NCT06116851), previously measured by the routine clinical electrochemiluminescence immunoassay (ECLIA; S-Testo, test code 2735) for testosterone, was analyzed using the developed LC–MS/MS assay. Then, for each sample, the percentage difference (Δ%) was calculated as the difference between concentrations measured by the two methods divided by their mean value. The difference was considered fitting the requirement when the percent difference was within ±20%

### ASSESSING THE PERFORMANCE OF THE METHOD IN FECAL SAMPLES

Fecal samples were prepared in triplicate from a suspension of fecal matter in EtOH at the ratio 1:8 (w/v). Two extraction methods were tested: MTBE extraction and extraction with a MeOH:water (80:20, v/v) mixture. Aliquots of the obtained suspensions (450 μL) were spiked with 20 μL of the IS solution and subjected to extraction with 1 mL of the corresponding extraction agent, according to the aforementioned extraction procedure for fecal samples.

Additionally, to evaluate the influence of sample weight on analytical performance, 3 sets of samples were prepared in triplicate containing 2 mg, 10 mg, and 50 mg of original fecal material. For the 2 mg and 10 mg sets, suspensions were prepared using ethanol at a 1:4 (w/v) ratio; for the 50 mg set, a 1:8 (w/v) ethanol suspension was used. Aliquots of 10 μL, 50 μL, or 450 μL of the corresponding suspensions were mixed with 20 μL of IS solution and extracted with 1 mL of MTBE according to the aforementioned extraction procedure for fecal samples.

### UHPLC-MS/MS ANALYSIS

A targeted UHPLC-MS/MS method was developed based on the previously published method^40^. The quantitative assay was expanded to enable simultaneous measurement of 21 steroids in a single run. To maximize MS sensitivity, the sMRM transition parameters were optimized for the instrument used. Analysis was performed on a SCIEX Triple Quad 7500 mass spectrometer, Sciex (Toronto, Canada), coupled with ExionLC, Sciex (Toronto, Canada). The equipment consisted of 2 ExionLC AD Pumps, ExionLC AD Autosampler, ExionLC AC Column Oven, and ExionLC System Controller. Chromatographic separation was performed using a Kinetex Biphenyl LC Column (100 × 2.1mm 1.7 μm), Phenomenex (Torrance, CA, USA), maintained at 35 °C. Injection volume was 10 μL. Separation was performed using gradient elution with 0.2 mM NH_4_F in water (v/v) (A) and 0.2 mM NH_4_F in water:MeOH (5:95, v:v) (B) at a flow rate of 0.3 mL/min. Gradient program was 0 min 40 % B, 3.50 min 68 % B, 9.5 min 71 % B, 13.5 min 80 % B, 14.5–19 min 100 % B, and period 19.1–21 min at 40 % B serving as equilibration time. Mass spectrometry was performed employing the OptiFlow Pro ion source, equipped with an analytical probe and E-lens. Data acquisition was performed in both negative and positive polarities with a capillary voltage of 1500 V in both modes. The source temperature was set to 400 °C. The nebulizer gas (GS1) and heater gas (GS2), both using air, were set to 40 psi and 70 psi, respectively. Curtain gas (nitrogen) was set to 40 psi. Analytes were detected using scheduled multiple reaction monitoring (Table S3).

## RESULTS AND DISCUSSION

### EVALUATION OF THE EXTRACTION METHODS IN PLASMA

To select the plasma sample preparation procedure, three extraction methods were evaluated. For that, we accessed the matrix-induced suppression by analysing the IS responses. The post-spiked IS signals were normalized to responses obtained at the same target concentrations in neat solutions. ( Figure **1**A and Table S4). The PP demonstrated the highest signal suppression for 9 steroids, while the SLE method demonstrated the highest suppression for 10 steroids. The LLE method provided the lowest suppression for 17 of the 19 steroids tested, with the exception of P4 and E2, for which the SLE method showed better performance. Overall, LLE exhibited the lowest matrix interference for the majority of analytes.

**Figure 1.**
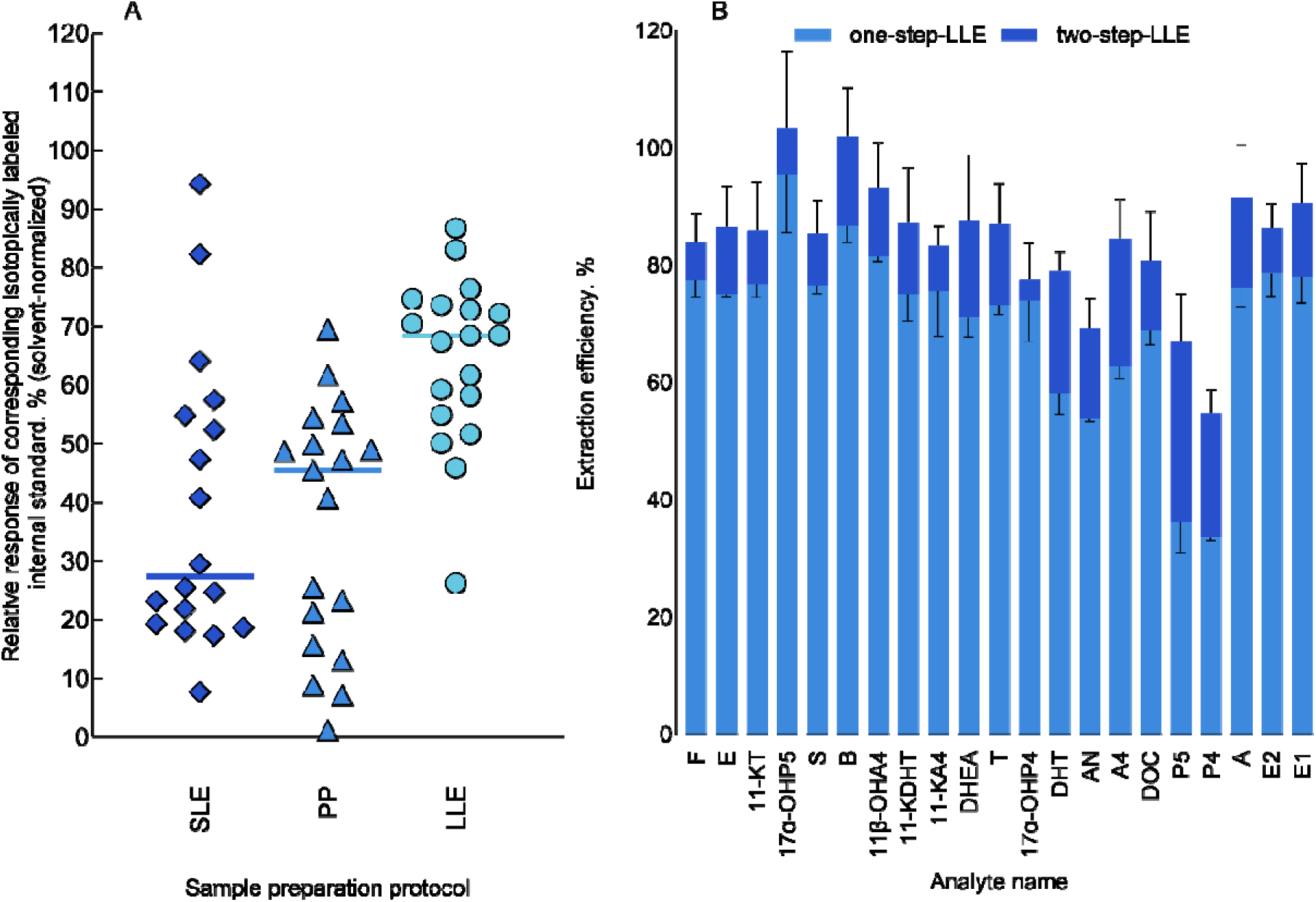
Steroid extraction tests. A - Comparison of the performance of three sampl preparation methods (PP, LLE, and SLE) for the steroid panel. The y-axis represents th normalized signal of the internal standard post-spiked into the extracted matrix and normalized to the same concentration signal in neat solutions. B - Comparison of sample preparation efficiencies using one-step LLE and two-step LLE for the steroid panel. Columns are grouped by analyte. Bar plots show mean extraction efficiency, calculated as the ratio of measured to nominal (expected) concentration, with error bars indicating SD. Concentrations were determined using a calibration curve prepared in neat solutions.

### COMPARISON OF ONE-STEP AND TWO-STEP LLE PROTOCOLS IN ADIPOSE TISSUE

Next, we sought to determine the distribution of the steroids between the aqueous and organic extraction phases during adipose tissue sample preparation. The calculated concentrations were used to evaluate the distribution of the steroids between the two extraction phases. The majority of the steroids showed extraction efficiencies between 50 and 100 in the methanol/water layer, whereas P4 and P5 showed significantly higher concentrations in the hexane layer (Figure S1). However, a second extraction step significantly improved the extraction efficiency for steroids that were poorly extracted in the first step ( Figure **1**B). For DHT, A4, P4, and P5, the extraction efficiency was improved by 20 percentage points when two-step-LLE was used. Therefore, two-step-LLE extractions were used for the rest of the adipose tissue extractions.

### LLOQ EVALUATION IN PLASMA, FECAL SAMPLES, AND ADIPOSE TISSUE

Due to the endogenous nature of the analytes, calibration curves were prepared in neat solutions. However, LLOQ values determined in neat solutions do not reflect actual quantification limits in biological matrices, as neat solutions are not subject to matrix effects or analyte losses during sample preparation. To address this limitation, LLOQ values were estimated using indirect approaches.

For plasma and fecal samples, analytical runs were evaluated (n=146 and n=151, respectively). LLOQ values obtained in neat solutions were converted to matrix-specific values using normalization based on IS response. In fecal samples homogenates 6 steroids were detected, and the LLOQ values ranged from 41.1 to 384 pM. In plasma, all the 21 analytes in the panel were detected with LLOQ values ranging from 12.0 to 1216 pM.

For adipose tissue, where a sufficiently large sample set was unavailable, a surrogate analyte approach was applied. LLOQ values for native steroids were estimated from the corresponding deuterated IS by applying transformation coefficients derived from signal-to-noise ratios. Using this approach, LLOQ values were successfully determined for 18 out of 21 steroids in the panel. The LLOQs ranged from 0.01 to 9 pmol/g. For most of the steroids, the LLOQ was below 1 pmol/g, except A, P5, AN, DHT, 17α-OHP5, and DHEA. The confirmed ULOQ levels were within the range of 200-2500 pmol/g (Tables 1 and S5–S7)

### MATRIX EFFECT IN ADIPOSE TISSUE

All analytes, except DHT, met the predefined acceptance criterion (CV < 15%), indicating the acceptability of the chosen method and the applied internal standards for the analysis of the assay mixture in human adipose tissue (Table S8). Further evaluation of DHT results showed that variability in the normalized matrix factor was caused by the variability in the internal standard signal, indicating the presence of matrix components that interfere with accurate determination of the internal standard and, therefore, accurate measurement of DHT concentration.

### INTRA-RUN REPEATABILITY, ACCURACY, AND PRECISION IN ADIPOSE TISSUE

Repeatability for each steroid was calculated as a CV between injections of each replicate (Figure S2). All the analytes with concentration clearly above the LLOQ met the set repeatability criteria. Increased variability in several cases was associated with signal proximity to the LLOQ (Table 1). The intra-run precision was acceptable for concentrations above the LLOQ (Table S9). The CV for F exceeded the acceptance criterion, possibly due to the uneven distribution of this steroid in adipose tissue. Next, we assessed the intra-run accuracy. Steroids in adipose tissue, apart from estrone (E1), showed acceptable precision and accuracy in our assay (Tables S10, S11). This indicates reduced method performance for estrone quantification, reflected in increased uncertainty in its measured concentrations.

**Table 1.** Determined analyte LLOQs in adipose tissue, fecal homogenates, and plasma samples.

| Analyte | Adipose tissue<br>LLOQ, pmol/g | Fecal<br>homogenate<br>LLOQ, pM | Plasma,<br>LLOQ, pM |
| --- | --- | --- | --- |
| DHEA | 8.9±0.7 | not detected | 837.8±149.1 |
| 11-KDHT | 0.52±0.06 | 116.6±30.3 | 25.4±3.7 |
| 11-KT | 0.1±0.01 | not detected | 12.9±1.5 |
| 17α-OHP5 | 6±0.2 | not detected | 1216.5±187.3 |
| T | 0.11±0.01 | 41.1±10.7 | 25.6±3.6 |
| 11β-OHA4 | 0.06±0.003 | not detected | 23.9±3.7 |
| AN | 3.8±0.2 | not detected | 137.9±29.4 |
| DHT | 5.7±0.2 | not detected | 414.4±94.5 |
| E1 | 0.12±0.01 | 166.1±97.5 | 12±1.9 |
| F | 0.28±0.01 | not detected | 49.3±8.5 |
| P5 | 3.8±0.2 | not detected | 1082.4±219.7 |
| DOC | 0.06±0.01 | not detected | 24.8±3.9 |
| S | 0.05±0.01 | not detected | 70.2±10.2 |
| 17α-OHP4 | 0.01±0.001 | not detected | 28.9±4.3 |
| A | 3.4±0.5 | not detected | 253.6±83.2 |
| E | 0.25±0.03 | not detected | 15.7±2.5 |
| P4 | 0.03±0.002 | 96.5±45.1 | 33±5.2 |
| A4 | 0.057±0.004 | 383.9±220.4 | 73±10.4 |
| E2 | not evaluated | 357.6±182 | 291.9±49 |
| B | not evaluated | not detected | 14.1±2.1 |
| 11-KA4 | not evaluated | not detected | 12.8±1.9 |

### INTER-RUN ACCURACY AND PRECISION USING SOLVENT- AND SERUM-BASED QC SAMPLES

Inter-run accuracy for solvent QC samples ranged between 100 and 112 %, and the precision was below 7.0% for all steroids present in the QC sample (Table S12-S15). Next, we sought to assess the reproducibility of our method by comparing T levels obtained using our method to those of a clinically validated ECLIA assay in 322 plasma samples. The calculated difference criterion was met for 231 of 322 (71.7%) samples. This shows a high degree of correlation between these 2 assays ( Figure **2**). Next, we measured steroids in commercially available serum quality control sets. MassCheck® Steroid Panel 1 included 5 steroids of the panel the MassTrak™ steroid serum QC set 1 included 10 (Table 2). The calculated accuracy for all compounds, except DHT, was within 85-115% at concentrations within the calibration range. However, DHT demonstrated substantial accuracy bias across all test levels (178%, 134%, and 155%), likely due to the matrix effect, which is not fully addressed in the current method. The method was considered suitable for DHT quantitation, although with a higher analytical error than for the other analytes.

**Figure 2.**
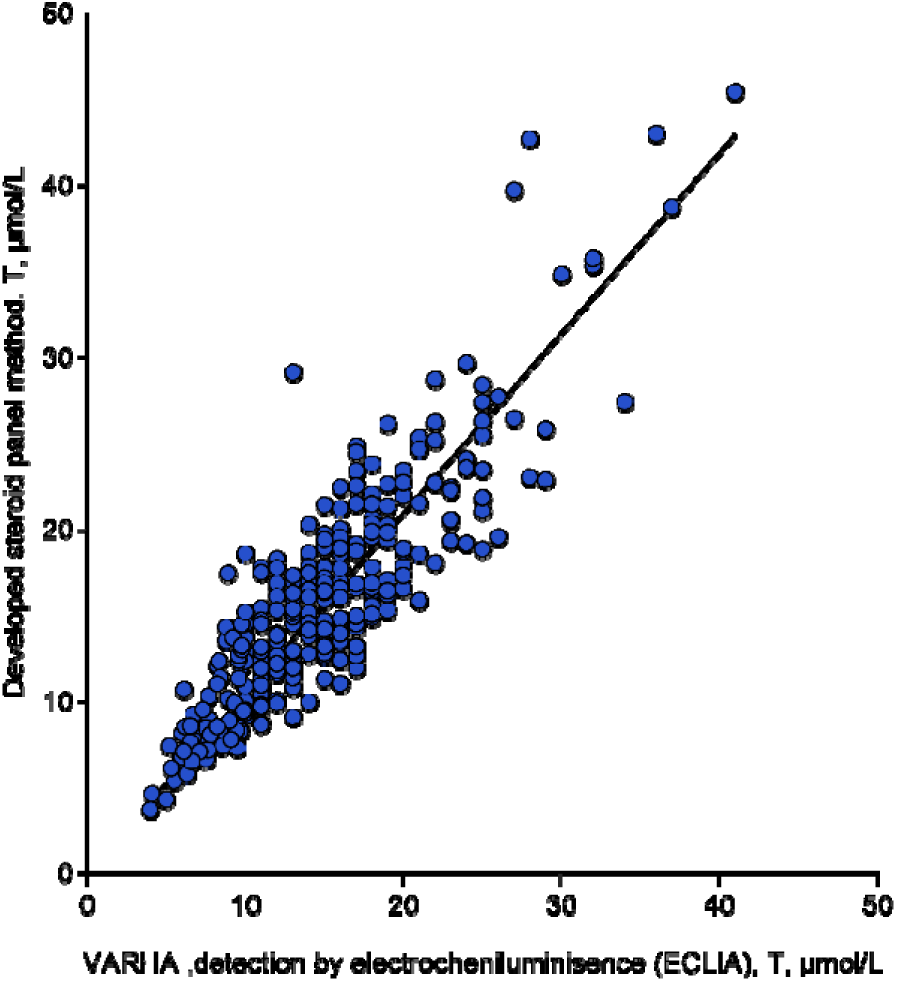
Comparing our UHPLC-MS/MS results to clinical chemistry assays. Correlation between testosterone concentrations measured by the UHPLC–MS/MS assay and those determined by a routine clinical chemistry assay in the accredited laboratory of the Wellbeing Services County of Southwest Finland (VARHA), R =0.880

**Table 2.** Intra-run accuracy and precision in the commercial serum control sets.

| Abbreviation | Conc. level | Nominal Concentration, pM | Measured concentration, pM | Accuracy, % | CV, % |
| --- | --- | --- | --- | --- | --- |
| P4 | Low† | 293 | 270 | 92 | 4.6 |
|  | Medium† | 2926 | 2582 | 88 | 1.2 |
|  | High*† | 97625 | 66176 | 68 | 13.1 |
| T | Low† | 225 | 199 | 88 | 3.1 |
|  | Medium† | 1730 | 1956 | 113 | 1.1 |
|  | High*† | 41604 | 43467 | 104 | 4.9 |
| A4 | Low† | 608 | 615 | 101 | 4.8 |
|  | Medium† | 6424 | 6383 | 99 | 3.3 |
|  | High*† | 109282 | 98966 | 91 | 7.0 |
| F | Low† | 28141 | 27363 | 97 | 3.3 |
|  | Medium*† | 271202 | 235894 | 87 | 3.0 |
|  | High*† | 886994 | 513585 | 58 | 13.2 |
| B | Low† | 514 | 479 | 93 | 6.9 |
|  | Medium† | 5051 | 4745 | 94 | 2.6 |
|  | High*† | 83413 | 72724 | 87 | 3.2 |
| S | Low† | 522 | 577 | 111 | 3.3 |
|  | Medium† | 5484 | 6161 | 112 | 0.4 |
|  | High*† | 94093 | 117408 | 125 | 6.0 |
| DOC | Low† | 115 | 107 | 93 | 4.6 |
|  | Medium† | 1047 | 1018 | 97 | 3.7 |
|  | High*† | 36616 | 37193 | 102 | 6.0 |
| 17α-OHP4 | Low† | 551 | 486 | 88 | 1.2 |
|  | Medium† | 5992 | 5219 | 87 | 1.1 |
|  | High*† | 188222 | 167777 | 89 | 5.2 |
| DHT | Low† | 358 | 639 | 178 | 4.3 |
|  | Medium† | 2823 | 3775 | 134 | 5.5 |
|  | High† | 5509 | 8560 | 155 | 5.8 |
| DHEA | Low† | 2659 | 2477 | 93 | 9.3 |
|  | Medium† | 25241 | 25782 | 102 | 6.8 |
|  | High† | 94999 | 108519 | 114 | 0.9 |
| A | Level I*‡ | 280 | 281 | 100 | 2.0 |
|  | Level II‡ | 685 | 773 | 113 | 0.5 |
|  | Level III‡ | 2620 | 2726 | 104 | 1.8 |
| B | Level I‡ | 2310 | 2130 | 92 | 0.7 |
|  | Level II‡ | 11500 | 10557 | 92 | 2.3 |
|  | Level III*‡ | 79700 | 68216 | 86 | 0.5 |
| F | Level I*‡ | 74500 | 69654 | 93 | 2.5 |
|  | Level II*‡ | 177000 | 151410 | 86 | 0.2 |
|  | Level III*‡ | 499000 | 304051 | 61 | 0.0 |
| E | Level I‡ | 5830 | 5670 | 97 | 2.2 |
|  | Level II*‡ | 34700 | 30752 | 89 | 0.9 |
|  | Level III*‡ | 85200 | 62104 | 73 | 2.4 |
| S | Level I‡ | 854 | 929 | 109 | 4.0 |
|  | Level II‡ | 4300 | 4700 | 109 | 0.3 |
|  | Level III‡ | 27800 | 28667 | 103 | 2.8 |

\*above upper limit of quantitation, † - Waters Masstrak™ Steroid Serum QC Set 1, ‡ - MassCheck® Steroid Panel 1 Serum Controls

### CARRYOVER EVALUATION USING UPPER-LEVEL CALIBRATION CURVE SAMPLES

We determined sample carryover by calculating the ratio of the analyte signal in a blank sample prepared in the solvent. The blank sample was injected immediately after an injection series of upper-level calibration curve samples (n=6). The signal elicited in this way was compared with the response of the samples in the calibration curve. Then the blank sample response was compared with the calibration curve. The calibration curve level providing 5-fold higher signal than in blank was considered as a concentration level not affected by carryover. The lowest calibration curve levels meeting the criteria are reported in the supplementary material (Table S16). The current settings are suitable for the vast majority of components. However, sample carryover was detected for A4, DOC,17_α_-OH-P4, P4 and T. Nevertheless, it should not significantly affect the analysis results, provided that the system is washed using a solvent blank injection after upper-level calibration curve injections and the samples are randomized and run within the same matrix. In that case, concentration variations are not dramatic and do not lead to significant errors in the results.

### ASSESSING THE PERFORMANCE OF THE METHOD IN FECAL SAMPLES

Extraction efficiency was initially assessed by comparing peak areas of isotopically labeled Internal standards ( Figure **3**A). With the exception of d4-E1, d8-E, d7-DOC, d4-DHT, and d7-S, MTBE showed a higher relative response than that observed with 80% MeOH in water, and MTBE extraction wa selected for further studies. The effect of sample weight on the analytical response was evaluated using normalized internal standard signals. We normalized the internal standard signal to that obtained with a 2 mg sample, since it contained the lowest amount of matrix. The resulting normalized values were multiplied by the weight-related expected signal enhancement factor: 1 for the 2 mg sample set, 5 for the 10 mg sample set, and 25 for the 50 mg sample set. The calculated response coefficient thus reflects the expected response ratio of native components. This takes into account the signal suppression by matrix components and the simultaneous signal enhancement due to the larger amount of analyte that underwent sample preparation. As shown in Figure **3**B, six analytes (d8-E, d3-11-KDHT, d4-DHT, d8-A, d4-E2, d4-E1) showed a significant increase in the response coefficient when a 50 mg sample weight was used. However, for most analytes, a sample weight of 10 mg yielded results that were better or comparable to those at 50 mg, while the 2 mg weight set was not optimal for any steroid in the panel. For subsequent experiments, we thus selected 10 mg as the optimal sample weight.

**Figure 3.**
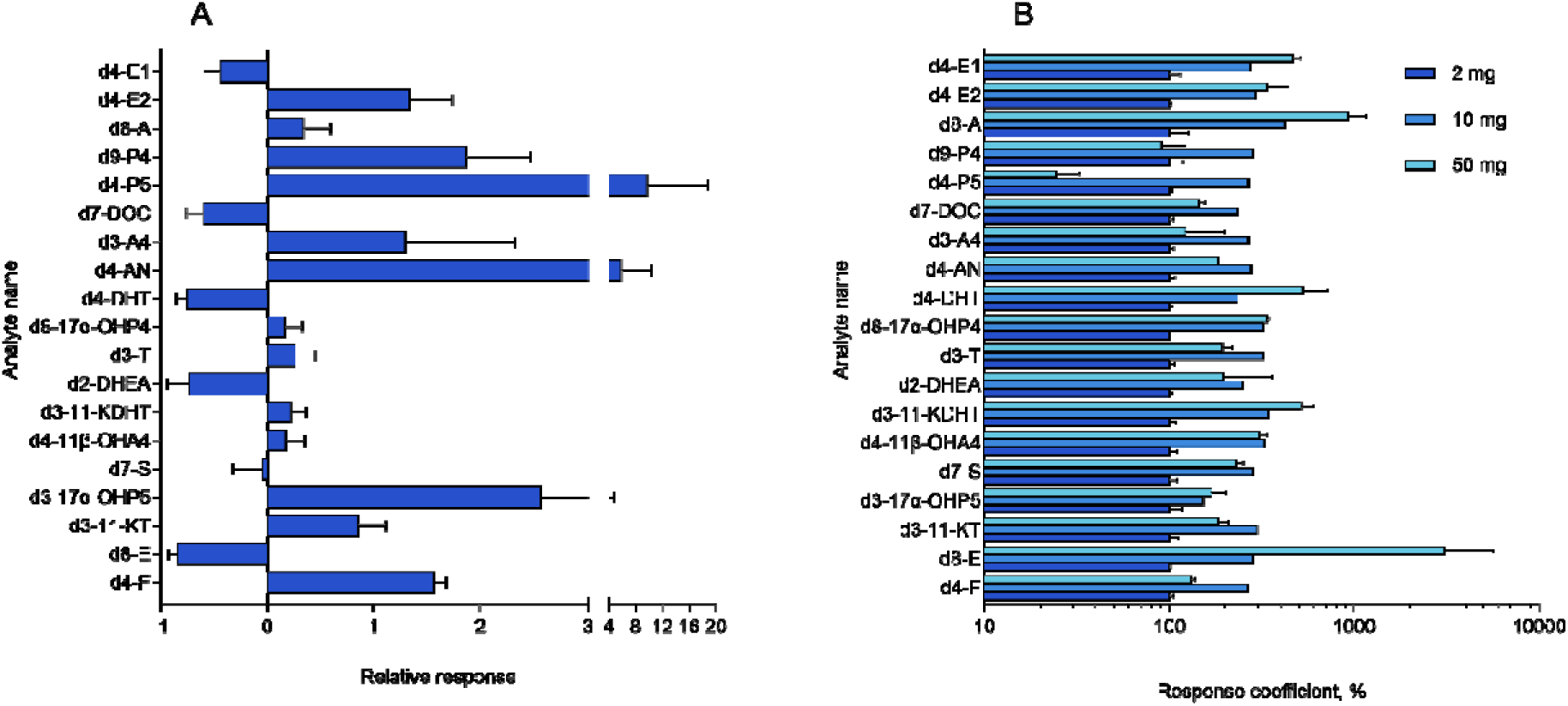
Optimization of the sample preparation conditions for fecal samples. A - Comparison of the mass spectrometer response for a selected panel of labeled steroid analytes after application of extraction methods with MTBE and 80% MeOH in water, expressed as relative response (MTBE extraction response/ 80%MeOH extraction response-1). Positive value indicate a benefit of MTBE extraction. B - Internal standard responses for sample preparation using 2, 10, and 50 mg sample weights. The responses were adjusted for a mass increase factor corresponding to the theoretically expected increase in component content and normalized to the lowest (2 mg) weight response. The lack of a proportional increase in response with increasing mass indicates the influence of matrix effect. Based on the combined results, 10 mg was selected as the optimal weight for the fecal samples preparation. In both panels (A and B), bar plots show mean values, with error bars indicating SD.

### STEROID LEVELS DETECTED IN ADIPOSE TISSUE OF WOMEN AND PLASMA, AND FECAL SAMPLES OF MEN

Replicate samples with different adipose tissue weights were prepared for analysis. The weights varied from 20 to 120 mg per replicate. Despite different initial weights, the results showed a high degree of data convergence in the range above the LLOQ. Based on the data obtained, concentrations of 4 analytes (A, AN, DHT, 11-KDHT) were below the LLOQ, concentrations of 3 analytes (E2, 11-KA4; B) were detected without LLOQ data available, and concentrations of the remaining 14 analytes were within the confirmed dynamic range ( Figure **4**A). We also measured steroid concentrations in plasma (Figure **4**B) and fecal samples ( Figure **4**C) from the PROMIC study (https://clinicaltrials.gov/study/NCT06116851) and were able to detect all 21 analytes in plasma and 11-KDHT, T, A4, P4, E2, and E1 in the fecal homogenates.

**Figure 4.**
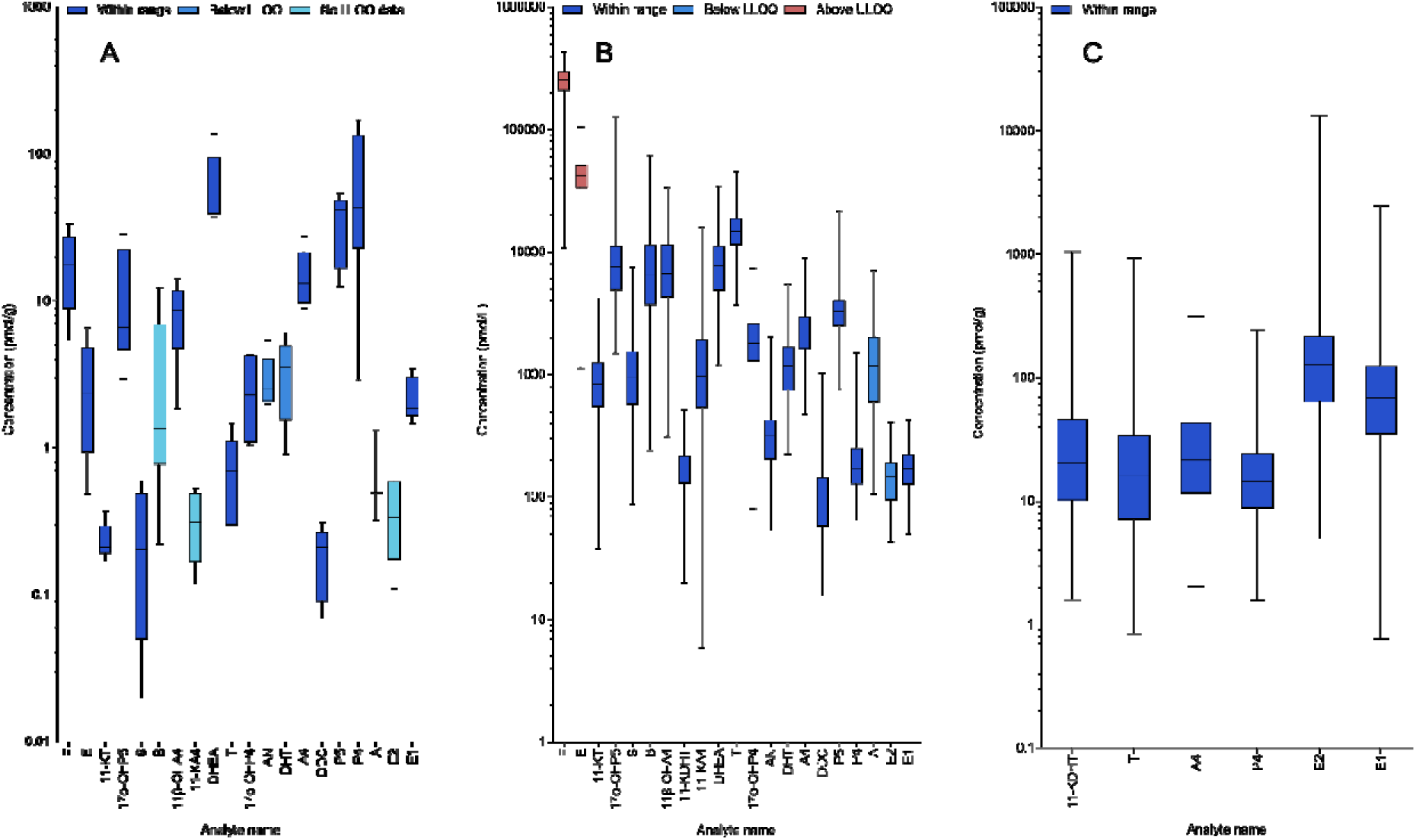
Validation in biological matrices. A – Adipose tissue samples from female donors (n = 15), obtained from 5 donors (3 samples per donor). 11-KDHT was not detected in the samples, B – blood plasma samples (n=325), male adult donors, C – fecal samples (n=332), male adult donors. In all panels, box-and-whisker plots show the concentration distribution, where the center line indicates the median, the box represents the interquartile range (25th–75th percentiles), and the whiskers extend to the minimum and maximum values.

### METHOD IMPLEMENTATION

We have applied the method to more than 600 samples in 8 separate studies within a year. The studies included analysis of different matrices such as plasma, fecal samples, and adipose tissue (Tables S17 and S18). The relative standard deviation for the solvent QC samples was <15% for all steroids except on one occasion for E1 (n=1, 15.4%). For the pooled QC samples the relative standard deviation were <25%, with a few exceptions, represented by single outliers for 17α-OHP5, 11-KDHT, P4, and E2

## CONCLUSIONS

The method described here enables quantification of an expanded steroid panel beyond routine clinical diagnostics, including the estrogens, androgens, progestogens, minerlacorticoids, and glucocorticoids, across complex matrices such as adipose tissue, fecal samples, plasma, and serum. Validation using isotope-labeled internal standards demonstrated robust quantitative performance in the presence of endogenous steroids. However, the quantification of E1 and DHT was associated with increased uncertainty, as E1 did not meet the acceptance criteria for accuracy and DHT showed significant matrix effects. The breadth of the panel permits a comprehensive assessment of steroid metabolism, while the achieved chromatographic resolution allows straightforward extension to additional metabolites.

## ACKNOWLEDGEMENTS

Mass spectrometry analysis was performed at the Turku Metabolomics Centre with the support of Biocenter Finland. The authors acknowledge the Turku Centre for Chemical and Molecular Analytics (CCMA), Turku, Finland, for providing equipment access for this study. This work was supported by the Doctoral Programme on Drug Research and Diagnostics (DRD), University of Turku Graduate School (UTUGS), University of Turku, Finland and by a grant from the Research Council of Finland (decision no. 347624) awarded to Dr. Alex Dickens.

Mass spectrometry analysis was performed at the Turku Metabolomics Centre with the support of Biocenter Finland. The authors acknowledge the Turku Centre for Chemical and Molecular Analytics (CCMA), Turku, Finland, for providing equipment access for this study.

## ASSOCIATED CONTENT

The following files are available free of charge.

Supporting_Information_Manuscript_Evstafev (PDF):

Figure S1. Distribution of steroids between two phases in LLE,

Figure S2. System repeatability. Distribution of coefficients of variation (CV) for each analyte,

Table S1. Abbreviations and purchase information of analytes and internal standards,

Table S2. Calibration curve levels, Table S3. LC-MS/MS conditions,

Table S4. Comparison of protein precipitation, LLE and SLE,

Table S5. Estimated LLOQ and ULOQ ranges for surrogate analytes and target analytes measured in adipose tissue,

Table S6. LLOQ evaluation in fecal samples, Table S7. LLOQ evaluation in plasma samples,

Table S8. Normalised matrix factors of steroids in human subcutaneous adipose tissue,

Table S9. Intra-run precision of each measured steroid in adipose tissue,

Table S10. Measured intra-run accuracy of the different steroids in adipose tissue without endogenous content correction,

Table S11. Measured intra-run accuracy in adipose tissue after removing the endogenous concentration,

Table S12. Intra-run precision and accuracy. Run 1,

Table S13. Intra-run precision and accuracy. Run 2,

Table S14. Intra-run precision and accuracy. Run 3,

Table S15. Inter-run precision and accuracy. Assessed on neat solutions quality control samples in plasma. Three separate run data is used,

Table S16. Degree of carryover observed,

Table S17. Pooled QC information for several studies

Table S18. Solvent QC information for several studies

## AUTHOR INFORMATION

### Author Contributions

The manuscript was written through contributions of all authors. All authors have given approval to the final version of the manuscript.

## ABBREVIATIONS

SD: standard deviation
LLOQ: lower limit of quantitation
ULOQ: upper limit of quantitation
QC: quality control
LLE: liquid-liquid extraction
PP: protein precipitation
SLE: supported liquid extraction

