## Supplemental Information for "Measurement of a panel of 21 steroids in a quantitative assay in human plasma, adipose tissue, and fecal samples using ultra-high-performance liquid chromatography–tandem mass spectrometry"

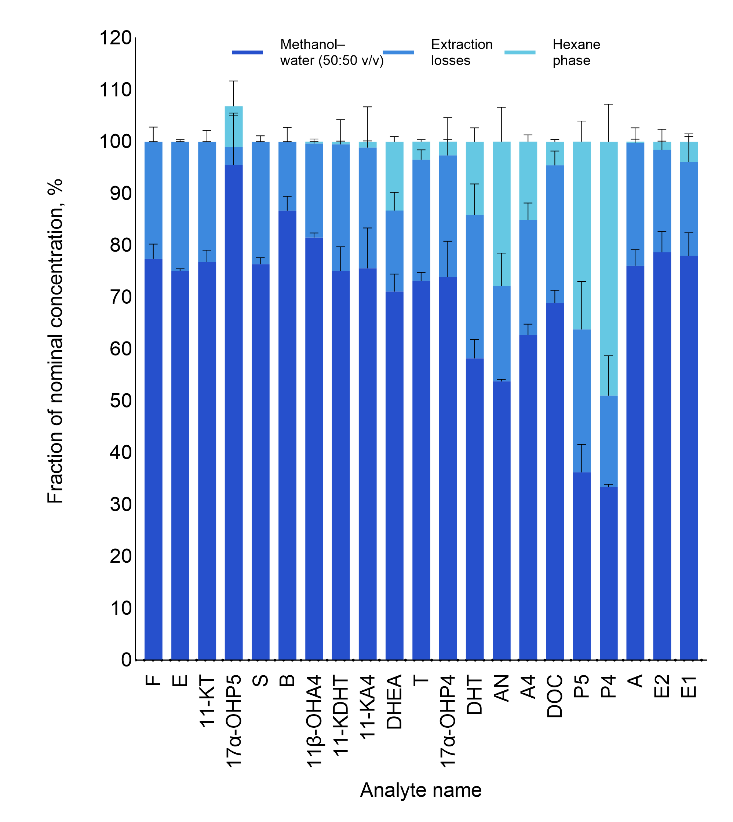

**Figure S1.** Distribution of steroids between two phases in LLE. The partitioned bars show the relative concentration in each phase relative to the total nominal concentration. The losses were calculated by subtraction of both layers’ concentration from the total nominal concentration. Concentrations were determined using a calibration curve prepared in neat solutions. Bar plots show mean values, with error bars indicating SD

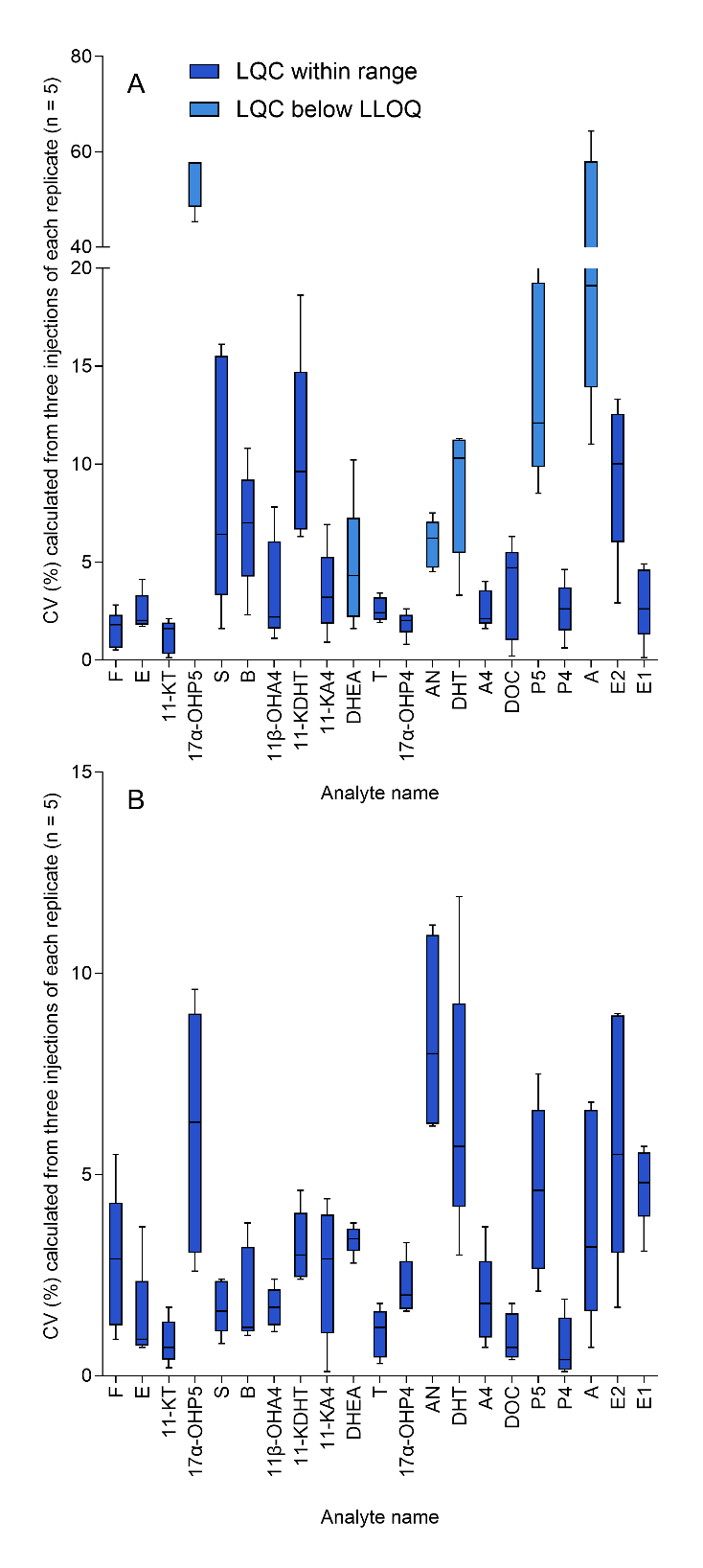

**Figure S2.** System repeatability. Distribution of coefficients of variation (CV) for each analyte. CV values are calculated based on all injections (n = 3) for each independent replicate (n = 5). High concentration quality control sample A – LQC samples, 1 nM; equivalent to 1 pmol/g in adipose tissue). B – HQC samples, 50 nM; equivalent to 50 pmol/g in adipose tissue). All samples were prepared from spiked adipose tissue homogenates. In all panels, box-and-whisker plots show the data distribution, where the center line indicates the median, the box represents the interquartile range (25th–75th percentiles), and the whiskers extend to the minimum and maximum values

Table S1. Abbreviations and purchase information of analytes and internal standards. NA= Not Available

| Abbreviation | IS/analyte | name | Other names | CAS | purity,% | part number | vendor |
| --- | --- | --- | --- | --- | --- | --- | --- |
| 11β-OHA4 | analyte | 11β-OH-Androstenedione | 4-androsten-11β-ol-3,17-dione, 11β-Hydroxyandrostenedione, 11-hydroxy-androstenedione | 382-44-5 | NA | 30410 | Cayman Chemical (Ann Arbor, MI, USA) |
| 11-KDHT | analyte | 11-Keto-Dihydrotestosterone | 5a-dihydro-11-ketotestosterone, 11-keto-dihydrotestosterone | 32694-37-4 | NA | 20200 | Cayman Chemical (Ann Arbor, MI, USA) |
| 11-KT | analyte | 11-ketotestosterone | 11-keto-testosterone | 564-35-2 | NA | 9002564 | Cayman Chemical (Ann Arbor, MI, USA) |
| AN | analyte | Androsterone | 3α-hydroxy-5α-androstan-17-one | 53-41-8 | 97 | [10601545](https://www.fishersci.fi/shop/products/androsterone-97-acros-organics-2/10601545) | Fisher scientific (Waltham, MA, USA) |
| P5 | analyte | Pregnenolone | pregn-5-en-3β-ol-20-one | 145-13-1 | 99 | 10582951 | Fisher scientific (Waltham, MA, USA) |
| DOC | analyte | 11-Deoxycorticosterone | 21-OH-Progesterone | 64-85-7 | 97 | D6875-500MG | Sigma-Aldrich (Saint Louis, MO, USA) |
| 17α-OHP4 | analyte | 17α-OH-Progesterone |  | 68-96-2 | NA | 46337-10MG | Sigma-Aldrich (Saint Louis, MO, USA) |
| 11-KA4 | analyte | Adrenosterone | 11-keto-androstenedione, 11-oxoandrostenedione, ndrost-4-ene-3,11,17-trione | 382-45-6 | 98 | 284998-5G | Sigma-Aldrich (Saint Louis, MO, USA) |
| A | analyte | Aldosterone |  | 52-39-1 | 95 | A9477-5MG | Sigma-Aldrich (Saint Louis, MO, USA) |
| A4 | analyte | Androstenedione | 4-androstenedione | 63-05-8 |  | 46033-250MG | Sigma-Aldrich (Saint Louis, MO, USA) |
| B | analyte | Corticosterone | 17-deoxycortisol, 11β,21-dihydroxyprogesterone | 50-22-6 | 98.5 | 27840-100MG | Sigma-Aldrich (Saint Louis, MO, USA) |
| F | analyte | Cortisol |  | 50-23-7 | certified reference material | C-106-1ML | Sigma-Aldrich (Saint Louis, MO, USA) |
| E | analyte | Cortisone |  | 53-06-5 | 98 | C2755-1G | Sigma-Aldrich (Saint Louis, MO, USA) |
| DHEA | analyte | Dehydroepiandrosterone | androstenolone | 53-43-0 | NA | NMID796B-10MG | Sigma-Aldrich (Saint Louis, MO, USA) |
| DHT | analyte | Dihydrotestosterone | 5α-dihydrotestosterone, 5α-DHT, androstanolone, stanolone | 521-18-6 | NA | NMID680-10MG | Sigma-Aldrich (Saint Louis, MO, USA) |
| E2 | analyte | Estradiol |  | 50-28-2 | NA | PHR1353-1G | Sigma-Aldrich (Saint Louis, MO, USA) |
| E1 | analyte | Estrone |  | 53-16-7 | 99 | E9750-1G | Sigma-Aldrich (Saint Louis, MO, USA) |
| P4 | analyte | Progesterone |  | 57-83-0 | 99 | P8783-1G | Sigma-Aldrich (Saint Louis, MO, USA) |
| T | analyte | Testosterone |  | 58-22-0 | 99 | 86500-1G | Sigma-Aldrich (Saint Louis, MO, USA) |
| S | analyte | 11-Deoxycortisol | cortodoxone, 17α,21-dihydroxyprogesterone, 17α,21-dihydroxypregn-4-ene-3,20-dione | 152-58-9 | 98 | D232600 | Toronto Research Chemicals (North York, ON, Canada) |
| 17α-OHP5 | analyte | 17α-OH-Pregnenolone | 17α-Hydroxypregnenolone, 17α OH-Pregnenolone, 17-OH-pregnenolone | 387-79-1 | 96 | H952320 | Toronto Research Chemicals (North York, ON, Canada) |
| d3-11-KT | IS | 11 ketotestosterone d3 |  |  |  | 9002754 | Cayman Chemical (Ann Arbor, MI, USA) |
| d3-11-KDHT | IS | 5α-dihydro-11-keto testosterone d3 |  |  |  | 9002761 | Cayman Chemical (Ann Arbor, MI, USA) |
| d3-T | IS | testosterone d3 |  | 77546-39-5 | 99.15 | 14037 | Isosciences (Ambler, PA, USA) |
| d7-A | IS | aldosterone d7 |  | 1261254-31-2 |  | S5093-0.1 | Isosciences (Ambler, PA, USA) |
| d4-DHT | IS | dihydrotestosterone d4 |  | 5295-66-9 | 99.24 | 15035 | Isosciences (Ambler, PA, USA) |
| d4-AN | IS | androsterone-d4 |  |  |  | NMID549-1MG | Sigma-Aldrich (Saint Louis, MO, USA) |
| d4-11β-OHA4 | IS | 11β-OH-4-androstene-3,17-dione-d4 |  |  |  | 903485-1MG | Sigma-Aldrich (Saint Louis, MO, USA) |
| d3-17α-OHP5 | IS | 17α-Hydroxy Pregnenolone-d3 |  |  | 97 | H952322 | Toronto Research Chemicals (North York, ON, Canada) |
| d8-B | IS | Corticosterone-d8 |  |  |  | C695702 | Toronto Research Chemicals (North York, ON, Canada) |
| d4-P5 | IS | Pregnenolone-d4 |  | 61574-54-7 |  | P712202 | Toronto Research Chemicals (North York, ON, Canada) |
| d7-S | IS | 11-Deoxy-Cortisol-d7 |  |  |  | D232602 | Toronto Research Chemicals (North York, ON, Canada) |
| d7-DOC | IS | 11-Deoxy-Corticosterone-d7 |  |  |  | D232593 | Toronto Research Chemicals (North York, ON, Canada) |
| d4-F | IS | Cortisol-d4 |  | 73565-87-4 |  | C696302 | Toronto Research Chemicals (North York, ON, Canada) |
| d9-P4 | IS | Progesterone-d9 |  | 15775-74-3 | 95 | P755902 | Toronto Research Chemicals (North York, ON, Canada) |
| d8-E | IS | Cortisone-d8 |  | N/A | 98 | C696502 | Toronto Research Chemicals (North York, ON, Canada) |
| d8-17α-OHP4 | IS | 17α-Hydroxy Progesterone-d8 |  | 850023-80-2 | 98 | H952332 | Toronto Research Chemicals (North York, ON, Canada) |
| d4-E2 | IS | 17b-Estradiol-d4 |  | 66789-03-5 | 97 | E888004 | Toronto Research Chemicals (North York, ON, Canada) |
| d4-E1 | IS | Estrone-d4 |  | 53866-34-5 | 97 | E889052 | Toronto Research Chemicals (North York, ON, Canada) |
| d2-DHEA | IS | d2 DHEA |  | 67034-83-7 | 98 | D229593 | Toronto Research Chemicals (North York, ON, Canada) |
| d3-A4 | IS | d3 Androstendione |  | 71995-66-9 | 97 | A637552 | Toronto Research Chemicals (North York, ON, Canada) |

Table S2. Calibration curve levels

| Level | Working solution, nM | Reconstituted LC-MS/MS sample (100 µL), pM | Calibration point conc. pM |
| --- | --- | --- | --- |
| L1 | 0.005 | 2 | 0.67 |
| L2 | 0.01 | 4 | 1.33 |
| L3 | 0.025 | 10 | 3.33 |
| L4 | 0.05 | 20 | 6.67 |
| L5 | 0.1 | 40 | 13.3 |
| L6 | 0.25 | 100 | 33.3 |
| L7 | 0.5 | 200 | 66.7 |
| L8 | 1 | 400 | 133 |
| L9 | 2.5 | 1000 | 333 |
| L10 | 5 | 2000 | 667 |
| L11 | 10 | 4000 | 1333 |
| L12 | 50 | 20000 | 6667 |
| L13 | 100 | 40000 | 13333 |
| L14 | 250 | 100000 | 33333 |
| L15 | 500 | 200000 | 66667 |
| L16 | 1000 | 400000 | 133333 |

Table S3. LC-MS/MS conditions

| Liquid chromatography conditions | | | | | | | | | | |
| --- | --- | --- | --- | --- | --- | --- | --- | --- | --- | --- |
| Instrumentation | | ExionLC AD Pump, AB3AD5976127 | | | | | | | | |
|  |  | ExionLC AD Pump, AB3AD5976128 | | | | | | | | |
|  |  | ExionLC AD Autosampler, AB3AC5972305 | | | | | | | | |
|  |  | AC Column Oven, AB2CT5972057 | | | | | | | | |
|  |  | System Controller, ABCBM5974024 | | | | | | | | |
|  |  | SCIEX Triple Quad™ 7500, FA220602109 | | | | | | | | |
| Chromatographic column | | Kinetex 1.7 um Biphenyl 100 A LC Column 100 x 2.1 mm (00D-4628-AN) | | | | | | | | |
| Phase A | | 0.2 mM NH_4_F in water | | | | | | | | |
| Phase B | | 0.2 mM NH_4_F in water/methanol (5/95) | | | | | | | | |
| Gradient | | time, min. | | | | Flow rate, µL/min | | %А | | %B |
|  |  | 0.0 | | | | 300 | | 60 | | 40 |
|  |  | 3.5 | | | | 300 | | 32 | | 68 |
|  |  | 9.5 | | | | 300 | | 29 | | 71 |
|  |  | 13.5 | | | | 300 | | 20 | | 80 |
|  |  | 14.5 | | | | 300 | | 0 | | 100 |
|  |  | 19.0 | | | | 300 | | 0 | | 100 |
|  |  | 19.1 | | | | 300 | | 60 | | 40 |
|  |  | 21.0 | | | | 300 | | 60 | | 40 |
| Column temperature | | 35 ºС | | | | | | | | |
| Autosampler temperature | | 10 ºС | | | | | | | | |
| Injection volume | | 10 µL | | | | | | | | |
| Analysis time | | 21 min | | | | | | | | |
| Source parameters | | | | | | | | | | |
| Ion source | | OptiFlow Pro | | | | | | | | |
| Ionization mode | | Positive (+Negative) | | | | | | | | |
| Source temperature, ºС | | 400 | | | | | | | | |
| Spray voltage, V | | 1500 | | | | | | | | |
| Ion source gas 1, psi | | 40 | | | | | | | | |
| Ion source gas 2, psi | | 70 | | | | | | | | |
| Entrance potential (EP), V | | 10 | | | | | | | | |
| MS/MS transitions conditions | | | | | | | | | | |
| Group | Compound ID | RT, min | RT window, (+/- s) | Q1 | Q2 | Dwell time, ms | CE, V | CXP, V | Q0 | ionization mode |
| F | d4-F | 5.4 | 45 | 367.3 | 121.1 | 128.3 | 29 | 16 | -10 | pos |
| F | F 1 | 5.4 | 45 | 363.1 | 121.2 | 128.3 | 31 | 8 | -10 | pos |
| F | F 2 | 5.4 | 45 | 363.1 | 91.0 | 128.3 | 83 | 10 | -10 | pos |
| E | d8-E | 6 | 45 | 369.2 | 169.0 | 70.8 | 33 | 20 | -10 | pos |
| E | E 1 | 6.1 | 45 | 361.1 | 163.1 | 70.8 | 31 | 26 | -10 | pos |
| E | E 2 | 6.1 | 45 | 361.1 | 77.0 | 70.8 | 107 | 10 | -10 | pos |
| 11-KT | d3-11-KT | 7.6 | 45 | 306.2 | 262.2 | 88.1 | 31 | 26 | -10 | pos |
| 11-KT | 11-KT 1 | 7.6 | 45 | 303.4 | 259.2 | 88.1 | 31 | 26 | -10 | pos |
| 11-KT | 11-KT 2 | 7.6 | 45 | 303.4 | 121.0 | 88.1 | 31 | 0 | -10 | pos |
| 17α-OHP5 | d3-17α-OHP5 | 7 | 45 | 336.2 | 300.2 | 55.5 | 15 | 17 | 10 | pos |
| 17α-OHP5 | 17α-OHP5 1 | 7 | 45 | 333.1 | 297.1 | 55.5 | 13 | 22 | 10 | pos |
| 17α-OHP5 | 17α-OHP5 2 | 7 | 45 | 333.5 | 315.1 | 55.5 | 10 | 18 | 10 | pos |
| S | d7-S | 8.6 | 45 | 354.1 | 100.1 | 3.0 | 33 | 16 | -10 | pos |
| S | S 1 | 8.8 | 45 | 347.1 | 97.0 | 3.0 | 27 | 12 | -10 | pos |
| S | S 2 | 8.8 | 45 | 347.1 | 109.0 | 3.0 | 33 | 16 | -10 | pos |
| B | d8-B | 9.6 | 45 | 355.5 | 337.0 | 26.5 | 32 | 11 | -10 | pos |
| B | B 1 | 9.6 | 45 | 347.5 | 329.2 | 26.5 | 21 | 22 | -10 | pos |
| B | B 2 | 9.6 | 45 | 347.1 | 121.1 | 26.5 | 29 | 8 | -10 | pos |
| 11β-OHA4 | d4-11β-OHA4 | 8.8 | 45 | 307.3 | 270.3 | 53.9 | 25 | 15 | -10 | pos |
| 11β-OHA4 | 11β-OHA4 1 | 8.9 | 45 | 303.2 | 267.2 | 53.9 | 24 | 14 | -10 | pos |
| 11β-OHA4 | 11β-OHA4 2 | 8.9 | 45 | 303.2 | 285.2 | 53.9 | 22 | 16 | -10 | pos |
| 11-KDHT | d3-11-KDHT | 8.9 | 45 | 308.4 | 254.2 | 100.0 | 25 | 24 | 50 | pos |
| 11-KDHT | 11-KDHT 1 | 8.9 | 45 | 305.4 | 251.2 | 100.0 | 25 | 21 | 50 | pos |
| 11-KDHT | 11-KDHT 2 | 8.9 | 45 | 305.4 | 287.2 | 53.9 | 23 | 20 | 50 | pos |
| 11-KA4 | 11-KA4 1 | 9.1 | 45 | 301.2 | 257.1 | 100.0 | 31 | 17 | -10 | pos |
| 11-KA4 | 11-KA4 2 | 9 | 45 | 301.2 | 121.0 | 61.0 | 31 | 15 | -10 | pos |
| DHEA | DHEA 1 | 10.3 | 45 | 289.1 | 253.0 | 43.2 | 15 | 46 | 30 | pos |
| DHEA | DHEA 2 | 10.3 | 45 | 289.4 | 271.1 | 43.2 | 11 | 28 | 30 | pos |
| DHEA | d2-DHEA 1 | 10.3 | 45 | 291.2 | 255.2 | 43.2 | 16 | 14 | 30 | pos |
| DHEA | d2-DHEA 2 | 10.3 | 45 | 291.2 | 273.2 | 43.2 | 12 | 15 | 30 | pos |
| T | d3-T | 11.6 | 45 | 292.1 | 97.0 | 62.9 | 29 | 12 | -10 | pos |
| T | T 1 | 11.7 | 45 | 289.1 | 97.0 | 62.9 | 29 | 12 | -10 | pos |
| T | T 2 | 11.7 | 45 | 289.1 | 109.2 | 62.9 | 31 | 6 | -10 | pos |
| 17αOH-P4 | d8-17αOH-P4 | 11.4 | 45 | 339.1 | 100.1 | 65.5 | 27 | 12 | -10 | pos |
| 17αOH-P4 | 17αOH-P4 1 | 11.5 | 45 | 331.1 | 109.1 | 65.5 | 31 | 12 | -10 | pos |
| 17αOH-P4 | 17αOH-P4 2 | 11.5 | 45 | 331.1 | 96.9 | 65.5 | 29 | 12 | -10 | pos |
| AN | d4-AN | 11.7 | 45 | 277.2 | 259.2 | 68.4 | 19 | 25 | -10 | pos |
| AN | AN 1 | 11.8 | 45 | 273.0 | 255.0 | 68.4 | 20 | 12 | -10 | pos |
| AN | AN 2 | 11.8 | 45 | 273.0 | 147.0 | 68.4 | 30 | 14 | -10 | pos |
| DHT | d4-DHT | 12.8 | 45 | 295.2 | 259.2 | 145.3 | 23 | 14 | 10 | pos |
| DHT | DHT 1 | 12.8 | 45 | 291.3 | 255.2 | 145.3 | 21 | 30 | 10 | pos |
| DHT | DHT 2 | 12.8 | 45 | 291.4 | 273.0 | 145.3 | 10 | 18 | 10 | pos |
| A4 | d3-A4 1 | 14.2 | 45 | 290.2 | 100.1 | 5.0 | 29 | 16 | -10 | pos |
| A4 | d3-A4 2 | 14.2 | 45 | 290.2 | 109.1 | 5.0 | 32 | 14 | -10 | pos |
| A4 | A4 1 | 14.3 | 45 | 287.0 | 97.0 | 5.0 | 27 | 14 | -10 | pos |
| A4 | A4 2 | 14.3 | 45 | 287.0 | 78.9 | 5.0 | 67 | 10 | -10 | pos |
| DOC | d7-DOC 1 | 14.7 | 45 | 338.2 | 100.1 | 92.9 | 28 | 11 | -10 | pos |
| DOC | d7-DOC 2 | 14.7 | 45 | 338.2 | 112.0 | 92.9 | 31 | 20 | -10 | pos |
| DOC | DOC 1 | 14.9 | 45 | 331.2 | 97.0 | 92.9 | 29 | 16 | -10 | pos |
| DOC | DOC 2 | 14.9 | 45 | 331.2 | 109.0 | 92.9 | 31 | 12 | -10 | pos |
| P5 | d4-P5 | 14.1 | 45 | 321.1 | 285.1 | 111.7 | 20 | 16 | -10 | pos |
| P5 | P5 1 | 14.2 | 45 | 317.5 | 299.1 | 111.7 | 15 | 16 | -10 | pos |
| P5 | P5 2 | 14.2 | 45 | 317.1 | 281.1 | 111.7 | 17 | 16 | -10 | pos |
| P4 | d9-P4 | 16 | 45 | 324.0 | 113.0 | 179.7 | 23 | 10 | -10 | pos |
| P4 | P4 1 | 16.1 | 45 | 315.1 | 97.1 | 179.7 | 23 | 10 | -10 | pos |
| P4 | P4 2 | 16.1 | 45 | 315.1 | 109.1 | 179.7 | 27 | 10 | -10 | pos |
| A | d8-A | 7.1 | 45 | 367.2 | 193.9 | 59.8 | -26 | -21 | 10 | neg |
| A | A 1 | 7.2 | 45 | 359.1 | 331.0 | 59.8 | -22 | -35 | 10 | neg |
| A | A 2 | 7.2 | 45 | 359.1 | 189.1 | 59.8 | -26 | -21 | 10 | neg |
| E2 | d4-E2 | 6.4 | 45 | 275.0 | 147.8 | 57.3 | -55 | -22 | 10 | neg |
| E2 | E2 1 | 6.4 | 45 | 271.0 | 182.9 | 57.3 | -52 | -19 | 10 | neg |
| E2 | E2 2 | 6.4 | 45 | 271.0 | 144.9 | 57.3 | -60 | -21 | 10 | neg |
| E1 | d4-E1 | 9.5 | 45 | 273.0 | 147.0 | 26.5 | -60 | -20 | 10 | neg |
| E1 | E1 1 | 9.6 | 45 | 269.1 | 144.9 | 26.5 | -48 | -15 | 10 | neg |
| E1 | E1 2 | 9.6 | 45 | 269.1 | 142.9 | 26.5 | -70 | -15 | 10 | neg |

Table S4. Comparison of protein precipitation, LLE and SLE. Response in matrix is normalized on response in neat solution (neat solution signal is considered 100%). The values shown belong to isotopically labeled internal standards

|  | F | E | 11-KT | 17α-OHP5 | S | B | 11β-OHA4 | 11-KDHT | 11-KA4 | DHEA | T | 17α-OHP4 | AN | DHT | A4 | DOC | P5 | P4 | A | E2 | E1 |
| --- | --- | --- | --- | --- | --- | --- | --- | --- | --- | --- | --- | --- | --- | --- | --- | --- | --- | --- | --- | --- | --- |
| neat solutions CV, % | 4.0 | 3.6 | 1.9 | 3.7 | 4.2 | 4.2 | 3.6 | 3.8 | 3.8 | 2.6 | 0.9 | 2.0 | 3.2 | 3.9 | 4.4 | 1.2 | 2.1 | 1.9 | 4.1 | 3.7 | 1.2 |
| SLE_IS normalised_1 | 7.95 | 17.99 | 30.77 | not detected | 18.42 | 18.42 | 26.78 | 18.85 | 18.85 | 24.87 | 56.43 | 42.55 | 24.03 | 57.80 | 52.45 | 64.93 | 27.66 | 49.18 | 83.26 | 92.70 | 18.25 |
| SLE_IS normalised_2 | 7.91 | 17.67 | 30.65 | not detected | 18.99 | 18.99 | 26.03 | 18.94 | 18.94 | 24.21 | 54.60 | 42.26 | 28.11 | 57.37 | 50.70 | 64.56 | 24.35 | 47.77 | 80.46 | 95.26 | 17.89 |
| SLE_IS normalised_3 | 7.00 | 16.45 | 26.89 | not detected | 16.88 | 16.88 | 23.61 | 18.20 | 18.20 | 20.44 | 53.39 | 37.55 | 13.42 | 57.24 | 54.02 | 62.85 | 22.13 | 45.17 | 83.00 | 95.06 | 21.85 |
| LLE_IS normalised_1 | 47.62 | 50.98 | 72.56 | 57.21 | 71.20 | 71.20 | 74.07 | 66.63 | 66.63 | 61.46 | 76.10 | 72.66 | 59.83 | 73.57 | 69.42 | 77.34 | 50.40 | 22.56 | 88.06 | 91.74 | 56.91 |
| LLE_IS normalised_2 | 48.20 | 51.29 | 70.71 | 54.28 | 67.56 | 67.56 | 71.82 | 65.81 | 65.81 | 62.09 | 72.60 | 67.43 | 56.31 | 71.80 | 69.05 | 74.17 | 41.54 | 31.57 | 79.24 | 78.91 | 57.25 |
| LLE_IS normalised_3 | 54.53 | 52.70 | 73.26 | 53.12 | 66.67 | 66.67 | 75.01 | 69.76 | 69.76 | 61.55 | 75.28 | 71.25 | 61.55 | 73.08 | 67.00 | 77.71 | 45.72 | 24.47 | 81.85 | 89.57 | 60.48 |
| PP_IS normalised_1 | 3.08 | 15.73 | 45.47 | 24.46 | 46.17 | 46.17 | 46.28 | 46.50 | 46.50 | 60.49 | 63.36 | 50.17 | 51.93 | 37.06 | 12.86 | 8.87 | 20.96 | 6.33 | 42.88 | 26.98 | 56.51 |
| PP_IS normalised_2 | 0.12 | 15.27 | 46.95 | 24.53 | 50.95 | 50.95 | 53.21 | 52.46 | 52.46 | 64.84 | 75.07 | 57.11 | 58.74 | 43.78 | 12.99 | 8.83 | 21.13 | 6.83 | 50.35 | 26.23 | 58.64 |
| PP_IS normalised_3 | 0.15 | 15.92 | 44.04 | 20.86 | 48.81 | 48.81 | 50.12 | 47.93 | 47.93 | 59.60 | 69.97 | 53.05 | 52.59 | 41.21 | 13.26 | 8.73 | 21.70 | 8.17 | 48.55 | 23.30 | 56.35 |
| mean SLE | 7.62 | 17.37 | 29.44 | 0.00 | 18.10 |  | 25.47 | 18.66 |  | 23.17 | 54.81 | 40.79 | 21.85 | 57.47 | 52.39 | 64.11 | 24.71 | 47.37 | 82.24 | 94.34 | 19.33 |
| mean LLE | 50.11 | 51.65 | 72.18 | 54.87 | 68.48 |  | 73.63 | 67.40 |  | 61.70 | 74.66 | 70.45 | 59.23 | 72.82 | 68.49 | 76.41 | 45.89 | 26.20 | 83.05 | 86.74 | 58.21 |
| mean PP | 1.12 | 15.64 | 45.48 | 23.28 | 48.64 |  | 49.87 | 48.97 |  | 61.64 | 69.46 | 53.44 | 54.42 | 40.68 | 13.04 | 8.81 | 21.27 | 7.11 | 47.26 | 25.50 | 57.16 |

Table S5. Estimated LLOQ and ULOQ ranges for surrogate analytes and target analytes measured in adipose tissue after following the two-step-LLE sample preparation procedure

| Surrogate analyte | Analyte | Transfor mation coefficient | Surrogate LLOQ, pmol/g | Surrogate ULOQ, pmol/g | Analyte LLOQ, pmol/g | Analyte ULOQ, pmol/g | Analyte LLOQ, pmol/g | Analyte LLOQ SD, pmols/g | Analyte ULOQ, pmol/g | Analyte ULOQ SD, pmols/g |
| --- | --- | --- | --- | --- | --- | --- | --- | --- | --- | --- |
| d2-DHEA | DHEA | 1.62 | 5 | 1000 | 8.12 | 1625 | 8.90 | 0.743 | 1780.31 | 148.7 |
|  |  | 1.92 |  |  | 9.60 | 1921 |  |  |  |  |
|  |  | 1.80 |  |  | 8.98 | 1796 |  |  |  |  |
| d3-11-KDHT | 11-KDHT | 1.18 | 0.5 | 500 | 0.59 | 589 | 0.52 | 0.064 | 519.62 | 63.9 |
|  |  | 1.01 |  |  | 0.51 | 507 |  |  |  |  |
|  |  | 0.93 |  |  | 0.46 | 463 |  |  |  |  |
| d3-11-KT | 11-KT | 0.90 | 0.1 | 250 | 0.09 | 226 | 0.10 | 0.0061 | 239.39 | 15.3 |
|  |  | 0.95 |  |  | 0.09 | 236 |  |  |  |  |
|  |  | 1.02 |  |  | 0.10 | 256 |  |  |  |  |
| d3-17α-OHP5 | 17α-OHP5 | 2.51 | 2.5 | 1000 | 6.28 | 2511 | 6.01 | 0.243 | 2404.52 | 97.4 |
|  |  | 2.38 |  |  | 5.96 | 2382 |  |  |  |  |
|  |  | 2.32 |  |  | 5.80 | 2320 |  |  |  |  |
| d3-T | T | 1.12 | 0.1 | 250 | 0.11 | 279 | 0.11 | 0.011 | 262.57 | 28.5 |
|  |  | 0.92 |  |  | 0.09 | 230 |  |  |  |  |
|  |  | 1.12 |  |  | 0.11 | 279 |  |  |  |  |
| d4-11β-OHA4 | 11β-OHA4 | 1.28 | 0.05 | 500 | 0.06 | 641 | 0.06 | 0.0033 | 631.68 | 32.5 |
|  |  | 1.19 |  |  | 0.06 | 596 |  |  |  |  |
|  |  | 1.32 |  |  | 0.07 | 659 |  |  |  |  |
| d4-AN | AN | 0.72 | 5 | 1000 | 3.61 | 723 | 3.81 | 0.173 | 762.06 | 34.5 |
|  |  | 0.79 |  |  | 3.94 | 788 |  |  |  |  |
|  |  | 0.78 |  |  | 3.88 | 775 |  |  |  |  |
| d4-DHT | DHT* | 2.32 | 2.5 | 1000 | 5.79 | 2317 | 5.73 | 0.227 | 2291.43 | 90.8 |
|  |  | 2.37 |  |  | 5.92 | 2367 |  |  |  |  |
|  |  | 2.19 |  |  | 5.48 | 2191 |  |  |  |  |
| d4-E1 | E1 | 1.26 | 0.1 | 500 | 0.13 | 632 | 0.12 | 0.012 | 601.06 | 60.3 |
|  |  | 1.06 |  |  | 0.11 | 532 |  |  |  |  |
|  |  | 1.28 |  |  | 0.13 | 640 |  |  |  |  |
| d4-F | F | 1.09 | 0.25 | 1000 | 0.27 | 1090 | 0.28 | 0.011 | 1139.05 | 45.9 |
|  |  | 1.18 |  |  | 0.30 | 1180 |  |  |  |  |
|  |  | 1.15 |  |  | 0.29 | 1147 |  |  |  |  |
| d4-P5 | P5 | 0.82 | 5 | 1000 | 4.08 | 817 | 3.84 | 0.240 | 768.68 | 48.0 |
|  |  | 0.77 |  |  | 3.84 | 768 |  |  |  |  |
|  |  | 0.72 |  |  | 3.60 | 721 |  |  |  |  |
| d7-DOC | DOC | 1.36 | 0.05 | 250 | 0.07 | 341 | 0.057 | 0.012 | 285.68 | 58.1 |
|  |  | 1.16 |  |  | 0.06 | 291 |  |  |  |  |
|  |  | 0.90 |  |  | 0.05 | 225 |  |  |  |  |
| d7-S | S | 0.78 | 0.05 | 250 | 0.04 | 196 | 0.048 | 0.0081 | 238.18 | 40.4 |
|  |  | 1.11 |  |  | 0.06 | 277 |  |  |  |  |
|  |  | 0.97 |  |  | 0.05 | 242 |  |  |  |  |
| d8-17α-OHP4 | 17α-OHP4 | 1.03 | 0.01 | 250 | 0.01 | 256 | 0.01 | 0.001 | 258.28 | 18.7 |
|  |  | 1.11 |  |  | 0.01 | 278 |  |  |  |  |
|  |  | 0.96 |  |  | 0.01 | 241 |  |  |  |  |
| d8-A | A | 3.98 | 1 | 100 | 3.98 | 398 | 3.37 | 0.535 | 336.72 | 53.5 |
|  |  | 3.11 |  |  | 3.11 | 311 |  |  |  |  |
|  |  | 3.01 |  |  | 3.01 | 301 |  |  |  |  |
| d8-E | E | 1.11 | 0.25 | 1000 | 0.28 | 1114 | 0.25 | 0.028 | 992.21 | 112.3 |
|  |  | 0.97 |  |  | 0.24 | 971 |  |  |  |  |
|  |  | 0.89 |  |  | 0.22 | 892 |  |  |  |  |
| d9-P4 | P4 | 0.61 | 0.05 | 500 | 0.03 | 304 | 0.030 | 0.0015 | 300.09 | 15.5 |
|  |  | 0.57 |  |  | 0.03 | 283 |  |  |  |  |
|  |  | 0.63 |  |  | 0.03 | 313 |  |  |  |  |
| d3-A4 | A4 | 1.05 | 0.05 | 250 | 0.05 | 263 | 0.057 | 0.0038 | 282.80 | 19.2 |
|  |  | 1.21 |  |  | 0.06 | 302 |  |  |  |  |
|  |  | 1.13 |  |  | 0.06 | 283 |  |  |  |  |

*For DHT, AN was used as the internal standard because, due to natural component impurity, signal intensity of the native analyte for high concentration was elevated and influenced accuracy. Transformation coefficient calculations were done using the d4-DHT/DHT combination similarly to other components

Table S6. LLOQ evaluation in fecal samples based on 151 male donor specimens analyzed. Shown numbers are for homogenate.

| Steroid analyte | 11-KDHT | T | A4 | P4 | E2 | E1 |
| --- | --- | --- | --- | --- | --- | --- |
| transition # | 11-KDHT 1 | T 1 | A4 1 | P4 1 | E2 1 | E1 1 |
| IS | d3-11-KDHT | d3-T | d3-A4 1 | d9-P4 | d4-E2 | d4-E1 |
| Neat solution LLOQ | 20.00 | 10.00 | 50.00 | 20.00 | 100.00 | 50.00 |
| Neat solution UlOQ | 20000 | 50000 | 50000 | 20000 | 50000 | 50000 |
| Internal standard AUC in neat solution samples. Used 64 solvent samples | | | | | | |
| Average of IS Area | 795026.617 | 16001893.7 | 11297176.1 | 18333221.8 | 155918.554 | 4257198.34 |
| StdDev of IS Area | 78096.9908 | 1447499.34 | 1099721.44 | 1587161.3 | 56638.3011 | 1903472.6 |
| Count of IS Area | 64 | 64 | 64 | 64 | 64 | 64 |
| CV, % | 9.8 | 9.0 | 9.7 | 8.7 | 36.3 | 44.7 |
| Internal standard AUC in matrix samples. Used 151 fecal samples | | | | | | |
| Average of IS Area | 1.36E+05 | 3.89E+06 | 1.47E+06 | 3.80E+06 | 4.36E+04 | 1.28E+06 |
| StdDev of IS Area | 3.28E+04 | 9.55E+05 | 8.32E+05 | 1.74E+06 | 1.55E+04 | 4.87E+05 |
| Count of IS Area | 151 | 151 | 151 | 151 | 151 | 151 |
| CV, % | 24.0 | 24.5 | 56.5 | 45.9 | 35.7 | 38.0 |
| Coefficient of LLOQ transformation is calculated as ratio of average IS response in matrix samples to solvent samples, SD is calculated according to the formula on the page | | | | | | |
| Coefficient | 0.17 | 0.24 | 0.13 | 0.21 | 0.28 | 0.30 |
| SD | 0.04 | 0.06 | 0.07 | 0.10 | 0.14 | 0.18 |
| CV coeff, % | 26.0 | 26.1 | 57.4 | 46.7 | 50.9 | 58.7 |
| matix LLOQ, pM | 116.6 | 41.1 | 383.9 | 96.5 | 357.6 | 166.1 |
| matrix ULOQ, pM | 116576 | 205431 | 383875 | 96533 | 178794 | 166120 |

**Table S7. LLOQ evaluation in plasma samples based on an analytical run of 146 male donor specimens.**

| Steroid analyte | F | E | 11-KT | 17α-OHP5 | S | B | 11β-OHA4 | 11-KDHT | 11-KA4 | DHEA | T | 17α-OHP4 | AN | DHT | A4 | DOC | P5 | P4 | A | E2 | E1 |
| --- | --- | --- | --- | --- | --- | --- | --- | --- | --- | --- | --- | --- | --- | --- | --- | --- | --- | --- | --- | --- | --- |
| transition # | F 1 | E 1 | 11-KT 1 | 17α-OHP5 1 | S 1 | B 1 | 11β-OHA4 1 | 11-KDHT 1 | 11-KA4 1 | DHEA 1 | T 1 | 17αOH-P4 1 | AN 1 | DHT 1 | A4 1 | DOC 1 | P5 1 | P4 1 | A 1 | E2 1 | E1 1 |
| IS | d4-F | d8-E | d3-11-KT | d3-17α-OHP5 | d7-S | d7-S | d4-11β-OHA4 | d3-11-KDHT | d3-11-KDHT | d2-DHEA 1 | d3-T | d8-17αOH-P4 | d4-AN | d4-DHT | d3-A4 1 | d7-DOC 1 | d4-P5 | d9-P4 | d8-A | d4-E2 | d4-E1 |
| Neat solution LLOQ | 13.3 | 6.7 | 6.7 | 333.3 | 33.3 | 6.7 | 13.3 | 13.3 | 6.7 | 333.3 | 13.3 | 13.3 | 66.7 | 133.3 | 33.3 | 13.3 | 333.3 | 13.3 | 133.3 | 133.3 | 6.7 |
| Neat solution UlOQ | 66666.7 | 33333.3 | 33333.3 | 133333.3 | 33333.3 | 33333.3 | 66666.7 | 33333.3 | 33333.3 | 133333.3 | 33333.3 | 13333.3 | 133333.3 | 133333.3 | 13333.3 | 13333.3 | 133333.3 | 13333.3 | 133333.3 | 133333.3 | 33333.3 |
| Internal standard AUC in neat solution samples. Used 42-52 solvent samples | | | | | | | | | | | | | | | | | | | | | |
| Average of IS Area | 2.14E+07 | 6.65E+06 | 4.54E+07 | 5.95E+06 | 6.27E+07 | 6.28E+07 | 2.89E+07 | 7.76E+05 | 7.83E+05 | 1.90E+07 | 1.47E+07 | 5.69E+07 | 1.33E+07 | 7.23E+07 | 1.04E+07 | 1.19E+08 | 1.43E+07 | 1.61E+07 | 4.59E+05 | 3.57E+05 | 9.74E+06 |
| StdDev of IS Area | 2.12E+06 | 5.24E+05 | 3.57E+06 | 4.71E+05 | 5.01E+06 | 5.31E+06 | 2.82E+06 | 6.28E+04 | 6.39E+04 | 1.38E+06 | 6.88E+05 | 4.21E+06 | 1.72E+06 | 9.38E+06 | 8.99E+05 | 1.13E+07 | 1.36E+06 | 9.15E+05 | 2.71E+04 | 2.30E+04 | 8.37E+05 |
| Count of IS Area | 52 | 51 | 52 | 43 | 43 | 52 | 52 | 47 | 47 | 43 | 49 | 46 | 45 | 45 | 43 | 46 | 43 | 46 | 42 | 46 | 50 |
| CV, % | 9.9 | 7.9 | 7.9 | 7.9 | 8.0 | 8.5 | 9.8 | 8.1 | 8.2 | 7.3 | 4.7 | 7.4 | 12.9 | 13.0 | 8.7 | 9.5 | 9.6 | 5.7 | 5.9 | 6.4 | 8.6 |
| Internal standard AUC in matrix samples. Used 146 plasma samples | | | | | | | | | | | | | | | | | | | | | |
| Average of IS Area | 5.80E+06 | 2.83E+06 | 2.34E+07 | 1.63E+06 | 2.98E+07 | 2.98E+07 | 1.61E+07 | 4.07E+05 | 4.07E+05 | 7.55E+06 | 7.67E+06 | 2.62E+07 | 6.40E+06 | 2.33E+07 | 4.74E+06 | 6.43E+07 | 4.39E+06 | 6.51E+06 | 2.41E+05 | 1.63E+05 | 5.42E+06 |
| StdDev of IS Area | 8.19E+05 | 3.98E+05 | 2.06E+06 | 2.15E+05 | 3.59E+06 | 3.59E+06 | 1.90E+06 | 5.02E+04 | 5.02E+04 | 1.23E+06 | 1.01E+06 | 3.40E+06 | 1.09E+06 | 4.36E+06 | 5.34E+05 | 8.13E+06 | 7.84E+05 | 9.64E+05 | 7.80E+04 | 2.53E+04 | 7.36E+05 |
| Count of IS Area | 146 | 146 | 146 | 146 | 146 | 146 | 146 | 146 | 146 | 146 | 146 | 146 | 146 | 146 | 146 | 146 | 146 | 146 | 146 | 146 | 146 |
| CV, % | 14.1 | 14.1 | 8.8 | 13.2 | 12.1 | 12.1 | 11.8 | 12.3 | 12.3 | 16.2 | 13.2 | 13.0 | 17.0 | 18.8 | 11.3 | 12.7 | 17.9 | 14.8 | 32.3 | 15.5 | 13.6 |
| Coefficient of LLOQ transformation is calculated as ratio of average IS response in matrix samples to solvent samples, SD is calculated according to the formula on the page | | | | | | | | | | | | | | | | | | | | | |
| Coefficient | 0.27 | 0.43 | 0.52 | 0.27 | 0.47 | 0.47 | 0.56 | 0.52 | 0.52 | 0.40 | 0.52 | 0.46 | 0.48 | 0.32 | 0.46 | 0.54 | 0.31 | 0.40 | 0.53 | 0.46 | 0.56 |
| SD | 0.05 | 0.07 | 0.06 | 0.04 | 0.07 | 0.07 | 0.09 | 0.08 | 0.08 | 0.07 | 0.07 | 0.07 | 0.10 | 0.07 | 0.06 | 0.09 | 0.06 | 0.06 | 0.17 | 0.08 | 0.09 |
| CV coeff | 17.2 | 16.1 | 11.8 | 15.4 | 14.5 | 14.7 | 15.3 | 14.7 | 14.8 | 17.8 | 14.0 | 14.9 | 21.3 | 22.8 | 14.2 | 15.8 | 20.3 | 15.9 | 32.8 | 16.8 | 16.1 |
| matix LLOQ | 49.3 | 15.7 | 12.9 | 1216.5 | 70.2 | 14.1 | 23.9 | 25.4 | 12.8 | 837.8 | 25.6 | 28.9 | 137.9 | 414.4 | 73.0 | 24.8 | 1082.4 | 33.0 | 253.6 | 291.9 | 12.0 |
| matrix ULOQ | 246521 | 78259 | 64611 | 486598 | 70241 | 70308 | 119542 | 63558 | 64134 | 335121 | 64089 | 28921 | 275882 | 414371 | 29214 | 24769 | 432950 | 33014 | 253551 | 291890 | 59935 |

Table S8. Normalised matrix factors of steroids in human subcutaneous adipose tissue

| Analyte | nMF1 | nMF2 | nMF3 | nMF4 | nMF5 | nMF6 | CV, % |
| --- | --- | --- | --- | --- | --- | --- | --- |
| F | 0.98 | 1.09 | 1.07 | 1.04 | 1.05 | 1.10 | 4.1 |
| E | 1.08 | 1.08 | 1.02 | 1.10 | 0.94 | 1.16 | 7.1 |
| 11-KT | 1.08 | 1.12 | 0.97 | 1.09 | 1.11 | 1.12 | 5.3 |
| 17α-OHP5 | 0.91 | 1.12 | 1.04 | 1.14 | 1.08 | 1.16 | 8.4 |
| S | 1.08 | 1.05 | 0.97 | 1.10 | 0.98 | 1.08 | 5.3 |
| B | 1.22 | 1.29 | 1.08 | 1.29 | 1.17 | 1.29 | 7.2 |
| 11β-OHA4 | 1.10 | 1.21 | 1.04 | 1.13 | 1.08 | 1.23 | 6.5 |
| 11-KDHT | 1.05 | 1.17 | 1.06 | 1.12 | 1.13 | 1.18 | 4.9 |
| 11-KA4 | 1.04 | 1.15 | 1.08 | 1.12 | 1.19 | 1.39 | 10.8 |
| DHEA | 1.13 | 1.17 | 0.99 | 1.05 | 1.06 | 1.10 | 5.8 |
| T | 1.05 | 1.13 | 0.98 | 1.09 | 1.09 | 1.06 | 4.8 |
| 17α-OHP4 | 1.04 | 1.10 | 1.02 | 1.05 | 1.04 | 0.99 | 3.5 |
| AN | 0.97 | 0.96 | 0.83 | 0.96 | 0.91 | 0.98 | 6.3 |
| DHT | 3.86 | 1.55 | 1.30 | 3.10 | 1.67 | 2.05 | 44.7 |
| A4 | 1.03 | 1.08 | 0.99 | 1.09 | 1.14 | 1.19 | 6.6 |
| DOC | 1.09 | 1.15 | 1.03 | 1.11 | 1.06 | 1.19 | 5.3 |
| P5 | 1.18 | 1.33 | 1.06 | 1.28 | 1.15 | 1.46 | 11.5 |
| P4 | 0.90 | 1.08 | 0.96 | 1.11 | 0.95 | 1.17 | 10.1 |
| A | 1.27 | 1.27 | 1.14 | 1.20 | 1.16 | 1.36 | 6.8 |
| E2 | 1.06 | 1.08 | 0.96 | 1.10 | 1.01 | 1.07 | 4.9 |
| E1 | 1.06 | 1.13 | 1.03 | 1.02 | 1.03 | 1.17 | 5.6 |

Table S9. Intra-run precision of each measured steroid using the CV% metric in adipose tissue

| Compound | within-run precision  LQC, CV, % | within-run precision  HQC, CV, % |
| --- | --- | --- |
| F | 32.8*** | 5.0 |
| E | 4.2 | 3.7 |
| 11-KT | 5.4 | 2.2 |
| 17α-OHP5 | N/A* | 5.0 |
| S | 7.2 | 2.4 |
| B | 8.3** | 3.9 |
| 11β-OHA4 | 10.5 | 1.0 |
| 11-KDHT | 10.1 | 2.6 |
| 11-KA4 | 5.6** | 2.8 |
| DHEA | 5.6* | 3.3 |
| T | 6.2 | 2.3 |
| 17α-OHP4 | 4.1 | 2.6 |
| AN | 8.1* | 7.6 |
| DHT | 15.4* | 3.6 |
| A4 | 2.3 | 2.0 |
| DOC | 7.8 | 1.3 |
| P5 | 8.9* | 5.9 |
| P4 | 5.6 | 1.7 |
| A | 15.2* | 10.1 |
| E2 | 8.1** | 1.8 |
| E1 | 4.6 | 2.0 |

*spiked concentration is below LLOQ, **LLOQ was not evaluated, ***variation between replicates could be caused by uneven distribution of F in adipose tissue

Table S10. Measured intra-run accuracy of the different steroids in adipose tissue without endogenous content correction

| Compound | Accuracy, % | | | | | intra-run accuracy |
| --- | --- | --- | --- | --- | --- | --- |
| F | 113.5 | 106.4 | 113.3 | 121.6 | 109.8 | 112.9 |
| E | 99.0 | 98.7 | 104.6 | 105.4 | 107.0 | 102.9 |
| 11-KT | 100.8 | 100.6 | 99.4 | 104.5 | 103.7 | 101.8 |
| 17α-OHP5 | 117.8 | 114.0 | 105.1 | 115.0 | 120.2 | 114.4 |
| S | 108.3 | 108.4 | 107.8 | 110.3 | 114.1 | 109.8 |
| B | 105.0 | 106.4 | 107.1 | 112.9 | 114.6 | 109.2 |
| 11β-OHA4 | 111.7 | 112.0 | 113.5 | 113.7 | 114.1 | 113.0 |
| 11-KDHT | 100.2 | 95.2 | 98.4 | 101.9 | 100.2 | 99.2 |
| 11-KA4 | 101.7 | 96.2 | 100.4 | 102.6 | 103.4 | 100.9 |
| DHEA | 184.5 | 178.6 | 190.0 | 188.3 | 176.0 | 183.5 |
| T | 108.7 | 103.9 | 105.6 | 109.2 | 109.3 | 107.3 |
| 17α-OHP4 | 102.4 | 102.2 | 103.4 | 107.5 | 107.5 | 104.6 |
| AN | 101.2 | 89.5 | 93.7 | 108.3 | 103.2 | 99.2 |
| DHT | 109.3 | 101.1 | 107.1 | 103.3 | 109.9 | 106.1 |
| A4 | 135.8 | 132.5 | 134.3 | 138.4 | 139.0 | 136.0 |
| DOC | 84.8 | 86.0 | 86.2 | 87.9 | 86.8 | 86.3 |
| P5 | 148.3 | 141.0 | 144.2 | 146.3 | 163.8 | 148.7 |
| P4 | 109.2 | 109.2 | 111.3 | 110.5 | 113.7 | 110.8 |
| A | 103.2 | 114.0 | 88.1 | 107.2 | 94.2 | 101.4 |
| E2 | 109.1 | 104.4 | 108.0 | 107.8 | 109.2 | 107.7 |
| E1 | 134.9 | 138.9 | 136.2 | 140.6 | 141.4 | 138.4 |

Table S11. Measured intra-run accuracy in adipose tissue after spiking a known concentration of each steroid. The accuracy was then determined after removing the endogenous concentration.

| Compound | Accuracy, % | | | | | intra-run accuracy |
| --- | --- | --- | --- | --- | --- | --- |
| F | 100.6 | 93.5 | 100.4 | 108.7 | 96.9 | 100.0 |
| E | 97.4 | 97.1 | 103.0 | 103.7 | 105.3 | 101.3 |
| 11-KT | 100.3 | 100.1 | 98.9 | 104.0 | 103.2 | 101.3 |
| 17α-OHP5 | 103.1 | 99.3 | 90.3 | 100.3 | 105.5 | 99.7 |
| S | 108.0 | 108.0 | 107.5 | 109.9 | 113.8 | 109.4 |
| B | 104.2 | 105.6 | 106.3 | 112.1 | 113.8 | 108.4 |
| 11β-OHA4 | 100.2 | 100.5 | 102.0 | 102.2 | 102.7 | 101.5 |
| 11-KDHT | 100.2 | 95.2 | 98.3 | 101.8 | 100.1 | 99.1 |
| 11-KA4 | 101.2 | 95.8 | 100.0 | 102.1 | 102.9 | 100.4 |
| DHEA | 111.8 | 105.8 | 117.3 | 115.6 | 103.2 | 110.7 |
| T | 107.1 | 102.3 | 104.0 | 107.6 | 107.7 | 105.7 |
| 17α-OHP4 | 97.7 | 97.5 | 98.8 | 102.9 | 102.8 | 99.9 |
| AN | 97.4 | 85.6 | 89.9 | 104.4 | 99.4 | 95.3 |
| DHT | 108.0 | 99.7 | 105.7 | 102.0 | 108.6 | 104.8 |
| A4 | 111.2 | 107.9 | 109.7 | 113.8 | 114.4 | 111.4 |
| DOC | 84.9 | 86.0 | 86.3 | 87.9 | 86.8 | 86.4 |
| P5 | 107.7 | 100.4 | 103.7 | 105.7 | 123.2 | 108.2 |
| P4 | 102.0 | 101.9 | 104.0 | 103.2 | 106.4 | 103.5 |
| A | 100.0 | 110.8 | 84.9 | 104.0 | 91.0 | 98.2 |
| E2 | 106.3 | 101.7 | 105.3 | 105.0 | 106.4 | 104.9 |
| E1 | 127.3 | 131.3 | 128.6 | 133.0 | 133.8 | 130.8 |

Table S12. Intra-run precision and accuracy. Assessed on neat solutions quality control samples in plasma. Run 1.

| Analyte | **F** | **E** | **11-KT** | **17α-OHP5** | **S** | **B** | **11β-OHA4** | **11-KDHT** | **11-KA4** | **DHEA** | **T** | **17α-OHP4** | **AN** | **DHT** | **A4** | **DOC** | **P5** | **P4** | **A** | **E2** | **E1** |
| --- | --- | --- | --- | --- | --- | --- | --- | --- | --- | --- | --- | --- | --- | --- | --- | --- | --- | --- | --- | --- | --- |
| Nominal conc., nM. | 20.0 | 20.0 | 20.0 | 20.0 | 20.0 | 20.0 | 20.0 | 20.0 | 20.0 | 20.0 | 20.0 | 20.0 | 20.0 | 20.0 | 20.0 | 20.0 | 20.0 | 20.0 | 20.0 | 20.0 | 20.0 |
| Measured concentration, nM. | 22.7 | 20.2 | 21.8 | 21.0 | 20.7 | 23.2 | 21.5 | 20.3 | 20.1 | 20.7 | 21.9 | 22.0 | 21.2 | 20.1 | 21.2 | 21.9 | 20.2 | 21.4 | 20.9 | 21.0 | 20.1 |
|  | 23.2 | 21.8 | 22.2 | 21.8 | 21.3 | 22.4 | 23.0 | 21.5 | 21.8 | 20.7 | 22.4 | 21.6 | 22.4 | 20.1 | 23.5 | 22.6 | 20.2 | 21.4 | 21.0 | 21.8 | 21.1 |
|  | 23.1 | 21.7 | 21.1 | 21.5 | 22.7 | 23.5 | 22.0 | 22.8 | 21.8 | 22.9 | 22.3 | 22.0 | 21.9 | 22.7 | 22.2 | 21.2 | 20.7 | 19.8 | 21.3 | 22.2 | 21.8 |
|  | 22.3 | 21.0 | 21.5 | 22.1 | 21.0 | 21.6 | 21.9 | 21.9 | 22.2 | 21.4 | 22.0 | 22.2 | 23.6 | 21.3 | 21.9 | 20.4 | 20.3 | 19.2 | 21.5 | 20.9 | 21.5 |
|  | 22.3 | 20.8 | 20.9 | 21.1 | 21.6 | 21.2 | 21.2 | 21.6 | 23.0 | 21.1 | 21.6 | 21.3 | 20.0 | 22.5 | 21.9 | 20.5 | 21.6 | 19.3 | 19.6 | 20.8 | 20.6 |
| Mean, nM. | 22.7 | 21.1 | 21.5 | 21.5 | 21.5 | 22.4 | 21.9 | 21.6 | 21.8 | 21.4 | 22.1 | 21.8 | 21.8 | 21.3 | 22.1 | 21.3 | 20.6 | 20.2 | 20.9 | 21.4 | 21.0 |
| SD, nM. | 0.42 | 0.65 | 0.54 | 0.45 | 0.77 | 0.97 | 0.70 | 0.89 | 1.08 | 0.90 | 0.31 | 0.36 | 1.33 | 1.27 | 0.86 | 0.95 | 0.60 | 1.11 | 0.75 | 0.63 | 0.67 |
| Accuracy, % | 113.6 | 105.6 | 107.5 | 107.6 | 107.3 | 111.9 | 109.5 | 108.1 | 109.0 | 106.8 | 110.3 | 109.0 | 109.0 | 106.7 | 110.7 | 106.6 | 103.0 | 101.0 | 104.4 | 106.8 | 105.1 |
| Precision (CV,%) | 1.87 | 3.07 | 2.51 | 2.09 | 3.59 | 4.32 | 3.19 | 4.11 | 4.93 | 4.21 | 1.40 | 1.67 | 6.11 | 5.93 | 3.88 | 4.44 | 2.91 | 5.50 | 3.59 | 2.96 | 3.20 |
| N | 5 | 5 | 5 | 5 | 5 | 5 | 5 | 5 | 5 | 5 | 5 | 5 | 5 | 5 | 5 | 5 | 5 | 5 | 5 | 5 | 5 |

Table S13. Intra-run precision and accuracy. Assessed on neat solutions quality control samples in plasma. Run 2.

| Analyte | **F** | **E** | **11-KT** | **17α-OHP5** | **S** | **B** | **11β-OHA4** | **11-KDHT** | **11-KA4** | **DHEA** | **T** | **17α-OHP4** | **AN** | **DHT** | **A4** | **DOC** | **P5** | **P4** | **A** | **E2** | **E1** |
| --- | --- | --- | --- | --- | --- | --- | --- | --- | --- | --- | --- | --- | --- | --- | --- | --- | --- | --- | --- | --- | --- |
| Nominal conc., nM. | 20.0 | 20.0 | 20.0 | 20.0 | 20.0 | 20.0 | 20.0 | 20.0 | 20.0 | 20.0 | 20.0 | 20.0 | 20.0 | 20.0 | 20.0 | 20.0 | 20.0 | 20.0 | 20.0 | 20.0 | 20.0 |
| Measured concentration, nM. | 21.6 | 20.7 | 21.9 | 20.9 | 22.1 | 23.3 | 21.3 | 22.7 | 22.8 | 20.7 | 23.4 | 22.2 | 23.7 | 21.9 | 22.9 | 22.7 | 21.7 | 20.8 | 21.5 | 21.8 | 21.7 |
|  | 21.7 | 20.4 | 21.0 | 20.9 | 21.4 | 21.5 | 20.7 | 21.7 | 21.7 | 21.2 | 22.3 | 20.8 | 20.7 | 21.8 | 22.4 | 21.4 | 19.4 | 20.2 | 19.5 | 19.7 | 20.0 |
|  | 21.3 | 19.9 | 21.2 | 19.4 | 22.5 | 20.2 | 21.0 | 21.5 | 22.4 | 20.7 | 21.7 | 21.6 | 21.9 | 21.1 | 21.5 | 21.0 | 20.0 | 18.0 | 20.2 | 21.0 | 21.2 |
|  | 21.4 | 19.6 | 21.4 | 19.8 | 21.3 | 21.3 | 21.3 | 20.6 | 21.4 | 21.3 | 21.4 | 21.0 | 22.4 | 21.4 | 19.9 | 21.6 | 18.8 | 17.9 | 19.4 | 20.3 | 20.1 |
|  | 21.2 | 20.0 | 20.6 | 21.3 | 21.1 | 20.1 | 20.2 | 20.8 | 22.0 | 20.9 | 21.3 | 20.6 | 21.6 | 21.8 | 22.8 | 20.5 | 20.0 | 18.1 | 19.2 | 20.2 | 20.7 |
|  | 21.7 | 18.7 | 20.6 | 19.0 | 20.5 | 18.7 | 20.3 | 22.0 | 22.8 | 19.7 | 22.2 | 20.6 | 21.7 | 21.6 | 21.5 | 20.7 | 18.8 | 18.8 | 20.0 | 20.9 | 20.3 |
| Mean, nM. | 21.5 | 19.9 | 21.1 | 20.2 | 21.5 | 20.8 | 20.8 | 21.6 | 22.2 | 20.7 | 22.0 | 21.1 | 22.0 | 21.6 | 21.8 | 21.3 | 19.8 | 19.0 | 20.0 | 20.6 | 20.7 |
| SD, nM. | 0.2 | 0.7 | 0.5 | 0.9 | 0.7 | 1.6 | 0.5 | 0.8 | 0.6 | 0.6 | 0.8 | 0.7 | 1.0 | 0.3 | 1.1 | 0.8 | 1.1 | 1.3 | 0.8 | 0.7 | 0.7 |
| Accuracy, % | 107 | 99.4 | 106 | 101 | 107 | 104 | 104 | 108 | 111 | 104 | 110 | 106 | 110 | 108 | 109 | 107 | 98.9 | 94.8 | 100 | 103 | 103 |
| Precision (CV,%) | 1.02 | 3.49 | 2.29 | 4.65 | 3.32 | 7.55 | 2.28 | 3.66 | 2.61 | 2.75 | 3.44 | 3.10 | 4.52 | 1.37 | 5.15 | 3.62 | 5.46 | 6.70 | 4.12 | 3.62 | 3.17 |
| N | 6 | 6 | 6 | 6 | 6 | 6 | 6 | 6 | 6 | 6 | 6 | 6 | 6 | 6 | 6 | 6 | 6 | 6 | 6 | 6 | 6 |

Table S14. Intra-run precision and accuracy. Assessed on neat solutions quality control samples in plasma. Run 3.

| Analyte | **F** | **E** | **11-KT** | **17α-OHP5** | **S** | **B** | **11β-OHA4** | **11-KDHT** | **11-KA4** | **DHEA** | **T** | **17α-OHP4** | **AN** | **DHT** | **A4** | **DOC** | **P5** | **P4** | **A** | **E2** | **E1** |
| --- | --- | --- | --- | --- | --- | --- | --- | --- | --- | --- | --- | --- | --- | --- | --- | --- | --- | --- | --- | --- | --- |
| Nominal conc., nM. | 20.0 | 20.0 | 20.0 | 20.0 | 20.0 | 20.0 | 20.0 | 20.0 | 20.0 | 20.0 | 20.0 | 20.0 | 20.0 | 20.0 | 20.0 | 20.0 | 20.0 | 20.0 | 20.0 | 20.0 | 20.0 |
| Measured concentration, nM. | 22.5 | 21.9 | 22.1 | 22.7 | 21.6 | 22.7 | 22.0 | 21.3 | 21.9 | 21.4 | 22.9 | 22.5 | 17.8 | 21.7 | 20.2 | 23.1 | 23.8 | 26.3 | 20.3 | 23.2 | 22.7 |
|  | 23.8 | 21.8 | 22.4 | 22.0 | 21.8 | 22.7 | 22.9 | 20.2 | 21.1 | 21.1 | 23.3 | 21.9 | 19.4 | 20.3 | 21.7 | 23.0 | 22.2 | 25.1 | 20.4 | 21.3 | 22.3 |
|  | 23.0 | 21.9 | 21.0 | 22.4 | 22.4 | 23.7 | 22.5 | 20.4 | 21.2 | 20.7 | 23.0 | 22.0 | 19.4 | 21.0 | 21.9 | 22.5 | 21.3 | 23.8 | 20.7 | 22.3 | 24.1 |
|  | 24.0 | 22.3 | 21.8 | 21.3 | 19.9 | 20.2 | 22.3 | 21.1 | 21.3 | 20.8 | 22.0 | 21.3 | 20.1 | 22.0 | 21.8 | 22.3 | 23.0 | 22.4 | 19.8 | 20.5 | 22.5 |
|  | 23.0 | 21.7 | 21.9 | 21.4 | 20.5 | 20.5 | 21.2 | 21.5 | 22.0 | 19.4 | 22.4 | 21.5 | 18.3 | 19.3 | 20.6 | 22.4 | 22.6 | 21.5 | 18.7 | 19.8 | 21.7 |
|  | 23.2 | 21.3 | 22.2 | 22.0 | 22.9 | 22.2 | 22.0 | 20.6 | 21.3 | 21.0 | 23.1 | 22.2 | 20.2 | 23.0 | 22.0 | 21.3 | 21.3 | 22.4 | 20.4 | 22.6 | 24.0 |
|  | 22.2 | 21.4 | 22.0 | 22.9 | 21.4 | 20.9 | 22.2 | 21.6 | 21.5 | 21.5 | 22.0 | 21.3 | 21.1 | 21.2 | 20.2 | 20.6 | 20.9 | 21.0 | 21.1 | 22.5 | 23.0 |
|  | 23.4 | 23.2 | 21.6 | 20.9 | 21.8 | 21.6 | 22.0 | 21.9 | 22.5 | 20.6 | 21.6 | 20.8 | 20.9 | 21.0 | 21.5 | 21.1 | 21.8 | 20.4 | 20.8 | 21.4 | 23.0 |
|  | 22.1 | 21.1 | 22.6 | 20.4 | 20.8 | 22.0 | 22.5 | 22.4 | 22.5 | 19.9 | 21.5 | 21.1 | 20.4 | 22.0 | 19.7 | 21.3 | 20.8 | 19.5 | 20.6 | 21.8 | 23.2 |
|  | 23.0 | 22.5 | 21.7 | 22.5 | 22.4 | 22.7 | 23.2 | 21.5 | 21.5 | 21.2 | 22.3 | 21.5 | 23.0 | 20.7 | 21.3 | 21.8 | 21.0 | 20.9 | 20.6 | 22.1 | 23.4 |
|  | 22.9 | 22.3 | 21.9 | 20.7 | 21.2 | 22.1 | 22.6 | 21.7 | 21.5 | 20.6 | 23.1 | 21.5 | 21.5 | 23.6 | 23.3 | 22.9 | 22.6 | 20.9 | 20.7 | 23.1 | 23.1 |
|  | 22.5 | 22.7 | 21.9 | 22.4 | 21.0 | 21.8 | 22.3 | 21.9 | 22.6 | 20.8 | 22.3 | 20.6 | 22.3 | 20.8 | 21.2 | 21.6 | 20.4 | 20.1 | 19.9 | 22.2 | 22.8 |
|  | 22.0 | 21.1 | 21.2 | 21.3 | 22.1 | 22.9 | 22.1 | 22.5 | 22.4 | 20.3 | 21.5 | 20.8 | 21.7 | 20.4 | 21.5 | 22.0 | 20.8 | 19.5 | 20.5 | 21.7 | 22.4 |
|  | 23.5 | 22.7 | 22.6 | 22.7 | 21.6 | 22.9 | 22.6 | 20.7 | 22.5 | 21.2 | 23.3 | 22.0 | 21.6 | 20.0 | 22.6 | 23.0 | 20.9 | 20.5 | 21.3 | 23.0 | 24.0 |
|  | 23.0 | 22.2 | 21.5 | 21.2 | 21.8 | 22.2 | 22.1 | 22.9 | 22.4 | 21.1 | 21.9 | 20.6 | 21.6 | 21.8 | 23.1 | 21.8 | 21.5 | 20.1 | 20.6 | 22.6 | 22.5 |
|  | 22.4 | 21.5 | 21.6 | 21.4 | 21.0 | 22.4 | 21.3 | 21.6 | 21.7 | 20.6 | 21.9 | 20.9 | 21.7 | 20.6 | 20.9 | 21.2 | 21.2 | 20.1 | 20.7 | 24.3 | 22.5 |
|  | 21.6 | 22.1 | 21.7 | 20.9 | 20.5 | 21.7 | 21.9 | 21.5 | 21.6 | 20.1 | 21.5 | 20.6 | 22.0 | 20.8 | 20.2 | 21.4 | 20.0 | 19.9 | 20.6 | 22.8 | 23.1 |
|  | 23.1 | 22.9 | 23.1 | 21.9 | 20.6 | 22.2 | 22.7 | 22.3 | 22.7 | 21.2 | 22.7 | 22.1 | 23.0 | 22.5 | 22.3 | 22.2 | 21.7 | 20.3 | 21.7 | 21.9 | 24.3 |
|  | 22.3 | 21.1 | 22.1 | 22.4 | 21.8 | 21.9 | 22.3 | 22.1 | 22.9 | 21.6 | 22.5 | 21.5 | 23.5 | 21.1 | 21.5 | 22.2 | 22.1 | 20.7 | 21.2 | 22.6 | 24.6 |
|  | 22.5 | 21.8 | 21.6 | 22.0 | 22.1 | 22.9 | 22.3 | 21.8 | 21.8 | 20.2 | 22.1 | 21.1 | 24.1 | 20.4 | 22.7 | 21.5 | 21.9 | 21.0 | 21.4 | 24.3 | 23.8 |
|  | 21.9 | 20.7 | 22.1 | 22.0 | 22.1 | 22.6 | 22.6 | 21.6 | 22.5 | 21.1 | 21.9 | 21.0 | 22.2 | 22.7 | 22.1 | 21.4 | 20.5 | 20.5 | 21.1 | 23.7 | 24.3 |
|  | 22.9 | 23.2 | 22.7 | 21.9 | 22.4 | 23.8 | 22.7 | 21.7 | 22.7 | 21.3 | 23.1 | 21.9 | 21.2 | 21.0 | 22.1 | 22.3 | 20.8 | 21.1 | 21.4 | 26.2 | 24.6 |
|  | 21.7 | 22.6 | 21.8 | 20.5 | 21.2 | 22.9 | 21.9 | 21.3 | 22.0 | 20.7 | 22.1 | 21.1 | 21.1 | 20.0 | 21.0 | 21.9 | 20.5 | 20.8 | 21.1 | 23.1 | 23.7 |
|  | 22.1 | 21.4 | 22.1 | 20.9 | 20.9 | 22.1 | 22.1 | 20.6 | 20.8 | 20.6 | 21.7 | 20.7 | 21.6 | 19.8 | 21.0 | 21.4 | 21.3 | 20.3 | 21.2 | 24.8 | 24.1 |
|  | 21.7 | 21.0 | 21.8 | 21.2 | 21.6 | 23.0 | 21.4 | 22.5 | 21.3 | 18.8 | 22.4 | 20.8 | 21.5 | 20.8 | 21.1 | 21.5 | 20.2 | 20.4 | 21.1 | 23.9 | 22.9 |
|  | 22.7 | 21.8 | 22.5 | 21.2 | 21.6 | 22.6 | 22.4 | 22.3 | 23.4 | 21.9 | 22.8 | 21.3 | 21.9 | 20.6 | 21.5 | 22.6 | 21.5 | 21.6 | 21.7 | 24.5 | 25.0 |
|  | 22.3 | 22.0 | 22.6 | 21.4 | 22.3 | 23.2 | 22.8 | 21.6 | 22.9 | 20.9 | 22.7 | 20.9 | 22.6 | 20.3 | 22.7 | 22.2 | 23.3 | 21.9 | 21.8 | 23.5 | 24.0 |
|  | 22.2 | 20.8 | 21.6 | 21.4 | 21.5 | 21.0 | 21.9 | 21.1 | 21.6 | 20.5 | 22.3 | 20.8 | 23.4 | 22.6 | 21.8 | 20.8 | 23.3 | 21.1 | 21.6 | 24.1 | 22.7 |
| Mean, nM. | 22.6 | 21.9 | 22.0 | 21.6 | 21.5 | 22.3 | 22.3 | 21.6 | 22.0 | 20.8 | 22.3 | 21.3 | 21.4 | 21.1 | 21.6 | 21.9 | 21.5 | 21.2 | 20.8 | 22.9 | 23.4 |
| SD, nM. | 0.63 | 0.70 | 0.48 | 0.71 | 0.70 | 0.86 | 0.46 | 0.67 | 0.66 | 0.67 | 0.57 | 0.55 | 1.49 | 1.06 | 0.90 | 0.68 | 1.02 | 1.59 | 0.66 | 1.37 | 0.84 |
| Accuracy, % | 113 | 109 | 110 | 108 | 108 | 111 | 111 | 108 | 110 | 104 | 112 | 106 | 107 | 106 | 108 | 110 | 108 | 106 | 104 | 114 | 116.8 |
| Precision (CV,%) | 2.80 | 3.22 | 2.17 | 3.30 | 3.27 | 3.87 | 2.06 | 3.11 | 3.00 | 3.20 | 2.54 | 2.56 | 6.96 | 4.99 | 4.19 | 3.09 | 4.75 | 7.50 | 3.17 | 5.97 | 3.59 |
| N | 28 | 28 | 28 | 28 | 28 | 28 | 28 | 28 | 28 | 28 | 28 | 28 | 28 | 28 | 28 | 28 | 28 | 28 | 28 | 28 | 28 |

Table S15. Inter-run precision and accuracy. Assessed on neat solutions quality control samples in plasma. Three separate run data is used

| Analyte | **F** | **E** | **11-KT** | **17α-OHP5** | **S** | **B** | **11β-OHA4** | **11-KDHT** | **11-KA4** | **DHEA** | **T** | **17α-OHP4** | **AN** | **DHT** | **A4** | **DOC** | **P5** | **P4** | **A** | **E2** | **E1** |
| --- | --- | --- | --- | --- | --- | --- | --- | --- | --- | --- | --- | --- | --- | --- | --- | --- | --- | --- | --- | --- | --- |
| Nominal conc., nM. | 20.0 | 20.0 | 20.0 | 20.0 | 20.0 | 20.0 | 20.0 | 20.0 | 20.0 | 20.0 | 20.0 | 20.0 | 20.0 | 20.0 | 20.0 | 20.0 | 20.0 | 20.0 | 20.0 | 20.0 | 20.0 |
| Measured concentration, nM. | 22.7 | 21.1 | 21.5 | 21.5 | 21.5 | 22.4 | 21.9 | 21.6 | 21.8 | 21.4 | 22.1 | 21.8 | 21.8 | 21.3 | 22.1 | 21.3 | 20.6 | 20.2 | 20.9 | 21.4 | 21.0 |
|  | 21.5 | 19.9 | 21.1 | 20.2 | 21.5 | 20.8 | 20.8 | 21.6 | 22.2 | 20.7 | 22.0 | 21.1 | 22.0 | 21.6 | 21.8 | 21.3 | 19.8 | 19.0 | 20.0 | 20.6 | 20.7 |
|  | 22.6 | 21.9 | 22.0 | 21.6 | 21.5 | 22.3 | 22.3 | 21.6 | 22.0 | 20.8 | 22.3 | 21.3 | 21.4 | 21.1 | 21.6 | 21.9 | 21.5 | 21.2 | 20.8 | 22.9 | 23.4 |
| Mean, nM. | 22.3 | 21.0 | 21.5 | 21.1 | 21.5 | 21.8 | 21.7 | 21.6 | 22.0 | 21.0 | 22.1 | 21.4 | 21.7 | 21.4 | 21.8 | 21.5 | 20.6 | 20.1 | 20.6 | 21.6 | 21.7 |
| SD, nM. | 0.684 | 1.014 | 0.435 | 0.773 | 0.035 | 0.864 | 0.755 | 0.026 | 0.197 | 0.356 | 0.173 | 0.350 | 0.312 | 0.222 | 0.293 | 0.335 | 0.882 | 1.130 | 0.519 | 1.129 | 1.467 |
| Accuracy, % | 111 | 105 | 108 | 106 | 107 | 109 | 108 | 108 | 110 | 105 | 111 | 107 | 109 | 107 | 109 | 108 | 103 | 101 | 103 | 108 | 108 |
| Precision (CV,%) | 3.1 | 4.8 | 2.0 | 3.7 | 0.2 | 4.0 | 3.5 | 0.1 | 0.9 | 1.7 | 0.8 | 1.6 | 1.4 | 1.0 | 1.3 | 1.6 | 4.3 | 5.6 | 2.5 | 5.2 | 6.8 |
| N | 39 | 39 | 39 | 39 | 39 | 39 | 39 | 39 | 39 | 39 | 39 | 39 | 39 | 39 | 39 | 39 | 39 | 39 | 39 | 39 | 39 |

Table S16. Degree of carryover observed in the subsequent blank injection when using 6 consequent upper-level calibration curve samples. The concentration presented in adipose tissue corresponding concentration units

| Analyte | The concentration could be quantified without interference from carryover, with the blank signal not exceeding 20% of the signal observed at the stated concentration (pmol/g). | Confirmed LLOQ level, pmol/g |
| --- | --- | --- |
| F | 0.25 | 0.28 |
| E | 0.25 | 0.25 |
| 11-KT | 0.25 | 0.1 |
| 17α-OHP5 | 10.0 | 6.0 |
| S | 0.25 | 0.05 |
| B | 0.25 | Not deteremined |
| 11β-OHA4 | 0.5 | 0.06 |
| 11-KDHT | 0.5 | 0.52 |
| 11-KA4 | 0.25 | Not deteremined |
| DHEA | 10 | 8.9 |
| T | 1 | 0.11 |
| 17α-OHP4 | 1 | 0.01 |
| AN | 0.5 | 3.8 |
| DHT | 2.5 | 5.7 |
| A4 | 1 | 0.057 |
| DOC | 1 | 0.06 |
| P5 | 2.5 | 3.8 |
| P4 | 2.5 | 0.03 |
| A | 5 | 3.4 |
| E2 | 2.5 | Not deteremined |
| E1 | 1 | 0.12 |

Table S17. Pooled QC information for several studies carried out within one year time. Orange = above ULOQ, yellow = below LLOQ (BQL), pink = outlier

| Study | matrix | sample prep | sample number | QC N. |  | F | E | 11-KT | 17α-OHP5 | S | B | 11β-OHA4 | 11-KDHT | 11-KA4 | DHEA | T | 17α-OHP4 | AN | DHT | A4 | DOC | P5 | P4 | A | E2 | E1 | date |
| --- | --- | --- | --- | --- | --- | --- | --- | --- | --- | --- | --- | --- | --- | --- | --- | --- | --- | --- | --- | --- | --- | --- | --- | --- | --- | --- | --- |
| 1 | human plasma | LLE | 106 | 14 | average, pM | 207279.7 | 38842.9 | 1017.7 | 2280.0 | 881.3 | 8702.6 | 8371.1 | 182.6 | 1545.2 | 9504.4 | 4431.1 | 1256.1 | BQL | 254.5 | 2358.5 | 71.4 | BQL | 1596.8 | 1471.0 | 235.6 | 345.8 | Nov.2022 |
|  |  |  |  |  | sd, pM | 6002.8 | 2368.7 | 134.2 | 331.6 | 41.7 | 314.0 | 186.1 | 78.8 | 65.0 | 687.5 | 90.1 | 45.7 | BQL | 41.2 | 71.3 | 11.5 | BQL | 59.4 | 78.4 | 18.1 | 29.3 |  |
|  |  |  |  |  | cv % | 2.9 | 6.1 | 13.2 | 14.5 | 4.7 | 3.6 | 2.2 | 43.1 | 4.2 | 7.2 | 2.0 | 3.6 | BQL | 16.2 | 3.0 | 16.0 | BQL | 3.7 | 5.3 | 7.7 | 8.5 |  |
| 2 | human adipose tissue homogenates | LLE | 6 | 3 | average, pM | 220.8 | 105.4 | 645.3 | BQL | BQL | 13.1 | 357.5 | BQL | 111075.8 | 843.5 | BQL | 88.9 | BQL | 2741.0 | 174.8 | BQL | BQL | 526.4 | BQL | BQL | BQL | Nov.2022 |
|  |  |  |  |  | sd, pM | 1.3 | 2.1 | 4.4 | BQL | BQL | 1.2 | 14.2 | BQL | 598.4 | 44.3 | BQL | 42.6 | BQL | 69.8 | 7.9 | BQL | BQL | 434.1 | BQL | BQL | BQL |  |
|  |  |  |  |  | cv, % | 0.6 | 2.0 | 0.7 | BQL | BQL | 9.3 | 4.0 | BQL | 0.5 | 5.2 | BQL | 47.9 | BQL | 2.5 | 4.5 | BQL | BQL | 82.5 | BQL | BQL | BQL |  |
| 3 | human plasma, QC control | LLE | 146 | 19 | average, pM | 283708.2 | 52707.8 | 1251.1 | 13283.6 | 1529.8 | 12163.3 | 11535.3 | 167.4 | 1809.2 | 13004.1 | 19065.7 | 2316.3 | 405.4 | 1702.7 | 2876.5 | 143.9 | 4140.1 | 274.5 | 843.4 | BQL | 228.2 | Jun.2023 |
|  |  |  |  |  | sd, pM | 14594.5 | 2113.9 | 26.8 | 1436.4 | 70.0 | 458.5 | 409.6 | 11.0 | 133.6 | 878.7 | 1185.6 | 64.5 | 60.7 | 130.6 | 98.9 | 14.3 | 599.2 | 13.1 | 70.5 | BQL | 20.3 |  |
|  |  |  |  |  | cv, % | 5.1 | 4.0 | 2.1 | 10.8 | 4.6 | 3.8 | 3.6 | 6.5 | 7.4 | 6.8 | 6.2 | 2.8 | 15.0 | 7.7 | 3.4 | 10.0 | 14.5 | 4.8 | 8.4 | BQL | 8.9 |  |
| 4 | human plasma, QC case | LLE | 146 | 19 | average, pM | 295437.8 | 48678.3 | 1252.5 | 14889.7 | 1961.4 | 14447.0 | 13131.1 | 202.0 | 1985.1 | 11977.3 | 19755.5 | 2547.6 | 394.4 | 1479.8 | 3157.3 | 164.9 | 3862.6 | 282.7 | 859.8 | BQL | 251.1 | Jun.2023 |
|  |  |  |  |  | sd, pM | 15689.1 | 2971.2 | 36.6 | 2038.1 | 117.5 | 723.6 | 315.4 | 13.0 | 120.4 | 816.8 | 854.2 | 124.7 | 56.5 | 122.2 | 147.0 | 14.4 | 540.5 | 17.7 | 97.6 | BQL | 20.7 |  |
|  |  |  |  |  | cv, % | 5.3 | 6.1 | 2.9 | 13.7 | 6.0 | 5.0 | 2.4 | 6.4 | 6.1 | 6.8 | 4.3 | 4.9 | 14.3 | 8.3 | 4.7 | 8.8 | 14.0 | 6.3 | 11.3 | BQL | 8.2 |  |
| 5 | human fecal samples.control | LLE | 150 | 19 | average, pM | BQL | 158.9 | 75.2 | 19399.1 | BQL | BQL | BQL | 685.8 | BQL | BQL | 654.6 | BQL | 6812.7 | 7844.8 | 765.0 | BQL | BQL | 441.1 | BQL | 4357.6 | 2934.8 | Aug.2023 |
|  |  |  |  |  | sd, pM | BQL | 23.4 | 7.3 | 4251.8 | BQL | BQL | BQL | 96.2 | BQL | BQL | 101.4 | BQL | 1466.9 | 1130.4 | 80.9 | BQL | BQL | 28.0 | BQL | 442.0 | 160.2 |  |
|  |  |  |  |  | cv, % | BQL | 14.7 | 9.7 | 21.9 | BQL | BQL | BQL | 14.0 | BQL | BQL | 15.5 | BQL | 21.5 | 14.4 | 10.6 | BQL | BQL | 6.4 | BQL | 10.1 | 5.5 |  |
| 6 | human fecal samples.case | LLE | 150 | 19 | average, pM | BQL | 149.9 | 93.8 | 33661.3 | BQL | BQL | BQL | 646.8 | BQL | BQL | 632.9 | BQL | 9645.1 | 9960.9 | 764.9 | BQL | BQL | 397.4 | BQL | 2520.3 | 2533.0 | Aug.2023 |
|  |  |  |  |  | sd, pM | BQL | 23.2 | 10.1 | 9500.3 | BQL | BQL | BQL | 55.3 | BQL | BQL | 110.5 | BQL | 2065.9 | 2022.5 | 87.4 | BQL | BQL | 15.8 | BQL | 240.6 | 163.8 |  |
|  |  |  |  |  | cv, % | BQL | 15.5 | 10.8 | 28.2 | BQL | BQL | BQL | 8.6 | BQL | BQL | 17.5 | BQL | 21.4 | 20.3 | 11.4 | BQL | BQL | 4.0 | BQL | 9.5 | 6.5 |  |
| 7 | human SC adipose tissue | 2LLE | 15 | 5 | average, pM | 8279.3 | 1186.0 | 98.7 | 5291.4 | 117.8 | 1701.4 | 3637.6 | BQL | 129.1 | 27648.7 | 297.8 | 1086.6 | 1084.9 | 682.2 | 6338.4 | 75.9 | 15803.5 | 31178.2 | 160.2 | 145.7 | 861.1 | Feb.2023 |
|  |  |  |  |  | sd, pM | 254.0 | 169.6 | 4.8 | 1069.2 | 4.2 | 137.5 | 66.7 | BQL | 9.4 | 814.3 | 4.6 | 46.3 | 28.7 | 92.0 | 332.0 | 4.9 | 1393.1 | 3870.1 | 26.5 | 20.1 | 30.7 |  |
|  |  |  |  |  | cv, % | 3 | 14 | 5 | 20 | 4 | 8 | 2 | BQL | 7 | 3 | 2 | 4 | 3 | 13 | 5 | 6 | 9 | 12 | 17 | 14 | 4 |  |
| 8 | human serum | LLE | 217 | 28 | average, pM | 256027.4 | 47041.0 | 1360.9 | 78859.9 | 1602.9 | 20568.8 | 13170.1 | 201.1 | 1408.9 | 6312.2 | 3178.9 | 1056.3 | BQL | 464.0 | 1853.2 | 197.6 | 1602.3 | 216.9 | 516.5 | 73.0 | 294.9 | Apr.2023 |
|  |  |  |  |  | sd, pM | 11073.9 | 1017.2 | 19.3 | 38953.0 | 59.1 | 604.5 | 226.0 | 7.4 | 75.5 | 365.8 | 63.3 | 20.4 | BQL | 54.9 | 72.3 | 12.1 | 366.8 | 23.9 | 56.5 | 19.5 | 33.0 |  |
|  |  |  |  |  | cv, % | 4.3 | 2.2 | 1.4 | 49.4 | 3.7 | 2.9 | 1.7 | 3.7 | 5.4 | 5.8 | 2.0 | 1.9 | BQL | 11.8 | 3.9 | 6.1 | 22.9 | 11.0 | 10.9 | 26.7 | 11.2 |  |

Table S18. Solvent QC information for several studies analyzed within a year time. pink = outlier

| project | matrix | samples N | STD N |  | F | E | 11-KT | 17α-OHP5 | S | B | 11β-OHA4 | 11-KDHT | 11-KA4 | DHEA | T | 17αOH-P4 | AN | DHT | A4 | DOC | P5 | P4 | A | E2 | E1 |
| --- | --- | --- | --- | --- | --- | --- | --- | --- | --- | --- | --- | --- | --- | --- | --- | --- | --- | --- | --- | --- | --- | --- | --- | --- | --- |
| 1 | solvent | 106 | 14 | average, pM | 10918.1 | 10391.6 | 10017.7 | 10059.5 | 10726.8 | 11289.5 | 10785.2 | 10469.4 | 10840.5 | 10247.2 | 10963.1 | 10868.1 | 11140.1 | 10244.3 | 10949.1 | 10929.4 | 10883.8 | 11073.3 | 9979.9 | 11210.8 | 11274.7 |
|  |  |  |  | accuracy, % | 109.2 | 103.9 | 100.2 | 100.6 | 107.3 | 112.9 | 107.9 | 104.7 | 108.4 | 102.5 | 109.6 | 108.7 | 111.4 | 102.4 | 109.5 | 109.3 | 108.8 | 110.7 | 99.8 | 112.1 | 112.7 |
|  |  |  |  | sd, pM | 183.2 | 380.2 | 1271.2 | 457.6 | 409.5 | 504.8 | 399.3 | 881.7 | 592.9 | 514.9 | 218.2 | 355.6 | 956.1 | 350.3 | 333.6 | 318.4 | 443.1 | 385.0 | 327.3 | 459.4 | 456.7 |
|  |  |  |  | cv, % | 1.7 | 3.7 | 12.7 | 4.5 | 3.8 | 4.5 | 3.7 | 8.4 | 5.5 | 5.0 | 2.0 | 3.3 | 8.6 | 3.4 | 3.0 | 2.9 | 4.1 | 3.5 | 3.3 | 4.1 | 4.1 |
| 2 | solvent | 146 | 19 | average, pM | 7353 | 7113 | 7039 | 6779 | 7299 | 7220 | 7176 | 7182 | 7212 | 6950 | 7164 | 7060 | 6893 | 7301 | 6871 | 7009 | 7014 | 7147 | 6716 | 7043 | 7043 |
|  |  |  |  | accuracy, % | 110.3 | 106.7 | 105.6 | 101.7 | 109.5 | 108.3 | 107.6 | 107.7 | 108.2 | 104.2 | 107.5 | 105.9 | 103.4 | 109.5 | 103.1 | 105.1 | 105.2 | 107.2 | 100.7 | 105.6 | 105.6 |
|  |  |  |  | sd, pM | 171 | 186 | 210 | 426 | 377 | 282 | 190 | 301 | 384 | 246 | 150 | 189 | 381 | 298 | 190 | 147 | 441 | 118 | 265 | 324 | 298 |
|  |  |  |  | cv, % | 2.3 | 2.6 | 3.0 | 6.3 | 5.2 | 3.9 | 2.6 | 4.2 | 5.3 | 3.5 | 2.1 | 2.7 | 5.5 | 4.1 | 2.8 | 2.1 | 6.3 | 1.6 | 3.9 | 4.6 | 4.2 |
| 3 | solvent | 150 | 19 | average, pM | 10921 | 10390 | 10577 | 10038 | 10696 | 11032 | 10734 | 10004 | 10844 | 9915 | 10660 | 10445 | 9525 | 10799 | 10517 | 10532 | 10368 | 9946 | 11030 | 11169 | 11556 |
|  |  |  |  | accuracy, % | 109.2 | 103.9 | 105.8 | 100.4 | 107.0 | 110.3 | 107.3 | 100.0 | 108.4 | 99.1 | 106.6 | 104.4 | 95.3 | 108.0 | 105.2 | 105.3 | 103.7 | 99.5 | 110.3 | 111.7 | 115.6 |
|  |  |  |  | sd, pM | 342 | 310 | 284 | 331 | 494 | 534 | 313 | 378 | 410 | 520 | 224 | 350 | 723 | 580 | 372 | 288 | 667 | 262 | 1207 | 1298 | 1774 |
|  |  |  |  | cv, % | 3.1 | 3.0 | 2.7 | 3.3 | 4.6 | 4.8 | 2.9 | 3.8 | 3.8 | 5.2 | 2.1 | 3.4 | 7.6 | 5.4 | 3.5 | 2.7 | 6.4 | 2.6 | 10.9 | 11.6 | 15.4 |
| 4 | solvent | 15 | 5 | average, pM | 7567.9 | 7034.2 | 7165.6 | 7174.4 | 7150.6 | 7455.4 | 7300.9 | 7202.7 | 7262.3 | 7120.6 | 7348.0 | 7263.9 | 7264.6 | 7112.2 | 7377.4 | 7103.0 | 6868.3 | 6732.2 | 6963.2 | 7118.6 | 7003.5 |
|  |  |  |  | accuracy, % | 113.5 | 105.5 | 107.5 | 107.6 | 107.3 | 111.8 | 109.5 | 108.0 | 108.9 | 106.8 | 110.2 | 109.0 | 109.0 | 106.7 | 110.7 | 106.5 | 103.0 | 101.0 | 104.4 | 106.8 | 105.0 |
|  |  |  |  | sd, pM | 141.4 | 216.1 | 179.7 | 149.8 | 256.9 | 321.8 | 232.9 | 295.9 | 358.3 | 299.8 | 103.1 | 121.4 | 444.0 | 422.0 | 286.3 | 315.4 | 199.9 | 370.4 | 250.1 | 210.5 | 224.3 |
|  |  |  |  | cv, % | 1.9 | 3.1 | 2.5 | 2.1 | 3.6 | 4.3 | 3.2 | 4.1 | 4.9 | 4.2 | 1.4 | 1.7 | 6.1 | 5.9 | 3.9 | 4.4 | 2.9 | 5.5 | 3.6 | 3.0 | 3.2 |
| 5 | solvent | 217 | 28 | average, pM | 7538.1 | 7291.4 | 7326.1 | 7215.0 | 7174.5 | 7417.4 | 7416.2 | 7192.3 | 7333.2 | 6918.9 | 7447.7 | 7095.1 | 7133.2 | 7049.4 | 7181.9 | 7297.1 | 7183.7 | 7069.4 | 6940.5 | 7617.8 | 7785.5 |
|  |  |  |  | accuracy, % | 113.1 | 109.4 | 109.9 | 108.2 | 107.6 | 111.3 | 111.2 | 107.9 | 110.0 | 103.8 | 111.7 | 106.4 | 107.0 | 105.7 | 107.7 | 109.5 | 107.7 | 106.0 | 104.1 | 114.3 | 116.8 |
|  |  |  |  | sd, pM | 211.2 | 234.7 | 158.8 | 238.2 | 234.7 | 287.2 | 152.8 | 223.3 | 219.9 | 221.7 | 189.1 | 182.0 | 496.5 | 352.0 | 301.0 | 225.4 | 341.4 | 530.3 | 219.8 | 454.9 | 279.8 |
|  |  |  |  | cv, % | 2.8 | 3.2 | 2.2 | 3.3 | 3.3 | 3.9 | 2.1 | 3.1 | 3.0 | 3.2 | 2.5 | 2.6 | 7.0 | 5.0 | 4.2 | 3.1 | 4.8 | 7.5 | 3.2 | 6.0 | 3.6 |
